# Long distance migration is a major factor driving local adaptation at continental scale in Coho Salmon

**DOI:** 10.1101/2021.11.22.469546

**Authors:** Quentin Rougemont, Amanda Xuereb, Xavier Dallaire, Jean-Sébastien Moore, Eric Normandeau, Eric B. Rondeau, Ruth E. Withler, Donald M. Van Doornik, Penelope A. Crane, Kerry A. Naish, John Carlos Garza, Terry D. Beacham, Ben F. Koop, Louis Bernatchez

## Abstract

Inferring the genomic basis of local adaptation is a long-standing goal of evolutionary biology. Beyond its fundamental evolutionary implications, such knowledge can guide conservation decisions for populations of conservation and management concern. Here, we investigated the genomic basis of local adaptation in the Coho salmon (*Oncorhynchus kisutch*) across its entire North American range. We hypothesized that extensive spatial variation in environmental conditions and the species’ homing behavior may promote the establishment of local adaptation. We genotyped 7,829 individuals representing 217 sampling locations at more than 100,000 high-quality RADseq loci to investigate how recombination might affect the detection of loci putatively under selection and took advantage of the precise description of the demographic history of the species from our previous work to draw accurate population genomic inferences about local adaptation. Results indicated that genetic differentiation scans and genetic-environment association analyses were both significantly affected by variation in recombination rate as low recombination regions displayed an increased number of outliers. By taking these confounding factors into consideration, we revealed that migration distance was the primary selective factor driving local adaptation and partial parallel divergence among distant populations. Moreover, we identified several candidates SNP associated with long distance migration and altitude including a gene known to be involved in adaptation to altitude in other species. The evolutionary implications of our findings are discussed along with conservation applications.

## Introduction

Both plant and animal biodiversity are currently declining at unprecedented rates due to human activities (Allendorf, 2017), reducing species’ capacity to retain genetic diversity, and their evolutionary potential. In this context, accurate understanding of a species’ demographic and genetic response to environmental change is of major importance for their management and conservation. In particular, spatially heterogeneous environments can impose different selective pressures that could lead to local adaptation of populations (Kawecki & Ebert, 2004). In aquatic ecosystems, there is ample geographic variation in environmental conditions among habitats occupied by different populations of a given species. These conditions include salinity, temperature gradients, and geological features (Grummer et al., 2019), all of which can favor spatially varying selection and contribute to local adaptation (Gagnaire, Normandeau, Côté, Møller Hansen, & Bernatchez, 2012; Micheletti, Matala, Matala, & Narum, 2018; Narum Shawn R., Di Genova Alex, Micheletti Steven J., & Maass Alejandro, 2018). Landscape genomics provides a valuable framework to study genome-wide adaptive variation and its interaction with ecological conditions (Grummer et al., 2019). Despite the growing evidence from landscape genomics for local adaptation associated with variation in ecological parameters in marine and aquatic habitats (e.g., (Cayuela et al., 2020; Gagnaire et al., 2012; Hecht, Thrower, Hale, Miller, & Nichols, 2012; Micheletti et al., 2018), the field remains dominated by research in terrestrial systems (Grummer et al., 2019).

In recent years, landscape genomic methods (Narum & Hess, 2011) and genotype-environment association analyses (GEA) (e.g.; Bernatchez, 2016; Forester, Lasky, Wagner, & Urban, 2018; Villemereuil, Frichot, Bazin, François, & Gaggiotti, 2014) have been developed to identify signals of adaptation. Many riverscape genomic studies have investigated the effect of temperature, precipitation regime, geology, various distance variables (elevation, migration distance) or barriers to migration (e.g. dams) on local adaptation (e.g. Moore et al. 2017; Michelleti et al. 2018, Dallaire et al. 2020). A major challenge in thoroughly interpreting genome scans and GEAs is to accurately take into account the effect of demographic history. In particular, vicariance events and spatio-temporal variation in population size and migration rate impact the distribution of genetic variation, primarily by distorting the site frequency spectrum (Galtier, Depaulis, & Barton, 2000). These effects are difficult to model in most genome scan methods (Lotterhos & Whitlock, 2015; Narum & Hess, 2011). This is particularly true for populations occupying aquatic habitats and distributed along fragmented landscapes, with variable rates of connectivity and variable effective population sizes (Bierne, Roze, & Welch, 2013; Fourcade, Chaput-Bardy, Secondi, Fleurant, & Lemaire, 2013). In particular, the signal of isolation by distance (IBD), along with a unidirectional expansion from a common source population, has been shown to confound genome scan results, leading toi increased rates of false positives (Battey, Ralph, & Kern, 2020; Meirmans, 2012). To complicate these inferences further, the biologically meaningful signal of selection may not readily be discernible from demographic stochasticity if adaptation is driven by small shifts in allele frequencies at multiple loci (i.e. polygenic selection, (Pritchard, Pickrell, & Coop, 2010), as opposed to adaptation driven by hard selective sweeps that leave more discernible patterns along the genome.

A second, often overlooked, challenge is the effect of variation in recombination rate on genome scans and GEAs (Lotterhos, 2019). Nucleotide diversity and genetic differentiation are known to vary with local variation in recombination rate along the genome, with higher differentiation observed in regions of lower recombination (Charlesworth, Morgan, & Charlesworth, 1993). A consequence for genome scans is an associated high false positive rate in regions of low recombination (Perrier & Charmantier, 2019) and, conversely, a high false negative rate in regions of high recombination (Booker, Yeaman, & Whitlock, 2020).

Here, we aim to elucidate the effects of these phenomena on inferences of local adaptation with the largest landscape genomic study in a non-model species to date by explicitly accounting for the confounding factors of historical demography and recombination rate. We focused on the Coho salmon (*Oncorynchus kisutch)*, a species that has suffered from significant demographic declines over decades (Gustafson et al., 2007; Oke et al., 2020) and which is of high conservation concern because of its importance for the recreational and indigenous subsistence fisheries it supports (Oke et al., 2020). Based on a small subset of samples used in the present study, we recently documented the demographic history of Coho salmon throughout its North American range (Rougemont et al., 2020). This study showed that most of the contemporary species’ genetic diversity originates from a single major refugium that was located in the southern part of the species range, and previously glaciated regions further north were colonized following postglacial demographic expansion from this refugium. Coho salmon are distributed from 37 to 67 degrees of latitude across a range of environments, from warm to cold temperatures (e.g., 16.3°C in California *versus* -2.32°C in Alaska, Worldclim database), with varying precipitation regimes (e.g., over 2,438 mm annually in the Haida Gwaii archipelago *versus* 848 mm in parts of Alaska), and across different geological rock types (Garrity & Soller, 2009). These environmental variables were shown to be involved in the local adaptation of a congeneric species, the Steelhead trout (*Oncorhynchus mykiss*) in part of its range (Michelleti et al., 2018). Moreover, the species exhibits a range of spawning migration distances, from a few kilometers in short coastal rivers to over 2000km into the Yukon River of Alaska (Sandercock, 1991). The combined effects of migratory distance and total elevation gain to the spawning site could constitute an important selective factor (Moore et al., 2017). Indeed, long-distance migration has an elevated bioenergetic cost and can drive local adaptation of morphological, physiological or behavioural traits for improved migration efficiency (Bernatchez & Dodson, 2011; Crossin et al., 2004; Eliason et al., 2011). Finally, the natal homing behavior of salmonids and relatively low dispersal between adjacent rivers (Quinn, 1984) should contribute to the development and maintenance of local adaptive differentiation between populations (Waples, Naish, & Primmer, 2020). Therefore, the spatial variation in environmental conditions and the species’ homing behavior likely promote the establishment of local adaptation in Coho salmon populations.

Here we assessed the respective roles of various environmental factors in shaping putative adaptive variation among Coho salmon populations throughout its North American range. Knowledge of the species’ demographic history enabled us to interpret the distribution of outlier regions in the genome more accurately (Rougemont et al. 2020). Given recent population divergence from a single source post glaciation, we hypothesized that if local adaptation is due to a few SNPs of large effect (e.g., a recent hard sweep), these signals should be widespread among all populations. In this case, we expect (i) large changes in allele frequencies at candidate SNPs, and (ii) similar changes at shared candidate SNPs or genomic regions across different geographic parts of the range where populations undergo similar challenges (e.g., similar challenges to migration to spawning grounds, or thermal stress), resulting in parallel changes in allele frequencies. Alternatively, local adaptation may proceed through polygenic selection (Yeaman, 2015). In this case we expect small, but co-varying allele frequency shifts at candidate SNPs. We take advantage of a large genomic dataset to: (i) document patterns of adaptive genomic variation and evaluate the occurrence of parallel allele frequency shifts at loci of strong effect across the species range ii) identify the most important environmental factors involved in shaping patterns of adaptive genomic variation (iii) investigate the potential confounding effect of recombination rate variation on the detection of putatively adaptive SNPs and explicitly account for this factor within differentiation-based analyses. Finally, we show the power of a recently developed machine learning method to easily investigate levels of population structure over such a large dataset.

## Methods

### Sampling, environmental data, and bioinformatics

A total of 7,829 Coho salmon individuals were collected from 215 freshwater locations along the Pacific coast of North America, from California to Alaska, and from one site in Russia **(Table S01, Fig 1, FigS01**). The median number of individuals per site was 38 [range 14-70]. Sampled individuals were distributed across a heterogeneous environment **(Table S01**), which we predict would result in a range of selective pressures across populations. We previously used data from approximately one quarter of the current samples to show that contemporary Coho salmon populations mostly radiated postglacially from a single major lineage. This system provides ideal conditions to understand how recent divergence favors or constrains parallel adaptation across space (Rougemont et al., 2020).

**Figure 1:**
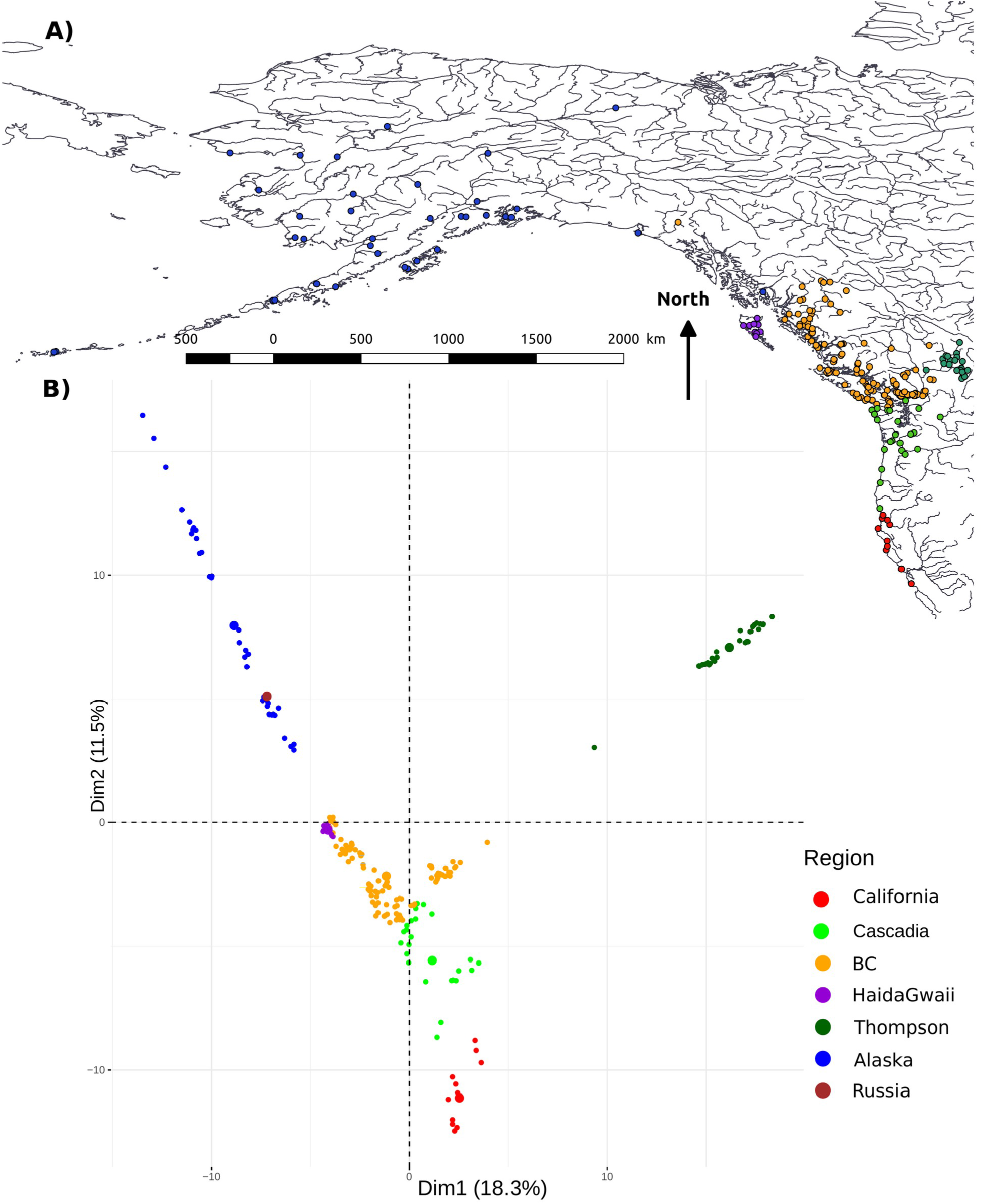
PCA genetic structure mirrors geography. A) Map of study area; dots represent samples and colors represent 7 major regional groups. B) PCA on allele frequency displaying the genetic structure among 215 Coho salmon populations grouped into the 7 different regions for illustrative purposes

Given the variability in temperature and precipitation across the study area, we delimited the catchment area upstream of the sampling sites and calculated the mean, minimum, maximum, range, and standard deviation of 19 climatic variables over these areas, resulting in 95 metrics associated with temperature (°C) and precipitation (mm) described in Supplementary Methods and provided in **Table S01 and Fig S02**. These were extracted from the WordclimV2.0 database (Fick & Hijmans, 2017) for the period 1970-2000. We reduced these data to a set of uncorrelated variables using two separate principal components analysis (PCA) and retained the significant axes of variation. The PCAs were performed using the R package ade4 (Dray & Dufour, 2007).We also extracted geological variables (rock type, geological era) from the United States Geological Survey database (Garrity & Soller, 2009), since geology has been identified as influencing population genetic structure in other salmonids (e.g., (Bourret, Dionne, Kent, Lien, & Bernatchez, 2013; Quéméré et al., 2016). Moreover, this parameter has been central to the designation of Evolutionary Significant Units in Coho salmon (Weitkamp et al., 1995).

Different Coho salmon populations undertake adult freshwater migrations from the marine environments to breeding sites that span a wide range of distances (river distances from <10 km to >2,300 km; **Table S01**), which we predicted should result in differential selective environments across populations (Olsen et al., 2011). In addition to distance to the breeding site, elevation is expected to exert a strong selective pressure on migration and homing phenotypes (e.g., Bernatchez & Dodson 1987). Such selection may result in differences in allele frequencies at loci linked to migration phenotypes associated with low or high elevation sites. Therefore, we computed the product of river length and altitude gain and standardized to a mean of zero and with a standard deviation of one (Moore et al. 2017). This measure will be referred to hereafter as «normalized distance». We evaluated the collinearity among predictor variables prior to our analysis using the variance inflation factor (VIF) and a correlation plot. No predictor displayed a VIF greater than 10, so all were retained for Genotype Environment Analyses (GEA).

DNA extractions and library preparation for ddRADseq were performed following protocols described elsewhere (Rougemont et al. 2020). Library preparation, PCR amplification and sequencing on the Ion Proton P1v2 chip were performed at the Genomic analysis platform of IBIS at Université Laval (http://www.ibis.ulaval.ca/). The bioinformatic procedure from Rougemont et al. (2020) was used, using Stacks v2 (Rochette, Rivera-Colón, & Catchen, 2019) and with the latest Coho salmon reference genome (https://www.ncbi.nlm.nih.gov/assembly/GCF_002021735.2/). The resulting vcf file was filtered using vcftools (Danecek et al., 2011). SNPs were retained if they displayed a read depth greater than 10 and less than 120. We then removed SNPs that were not present in at least 92% of individuals in the dataset. For each RAD locus, the single SNP with the highest minor allele frequency (MAF) was retained. More details about the laboratory protocol and bioinformatic procedure along with a table of the different steps can be found in supplementary methods and **Table S02**. This filtered dataset contained 105,362 SNPs.

### Population genetics and demography

We previously observed a clinal distribution of decreasing genetic diversity from the South to the North of the Pacific Coast. Using demographic modeling we resolved the species historical demography and showed that it has expanded northward from a single major southern refugium, resulting in an increase of the deleterious load due to surfing of deleterious mutations (Rougemont et al., 2020). We seek to verify whether our previous observations that (i) genetic diversity decreased northwards due to population expansion from a major southern refugium, and (ii) genetic differentiation increased toward the north, could be replicated by increasing the size from our previously published dataset from 58 populations to the full set of 217 sampling localities presented here.

As previously, we estimated gene diversity (Hs) and population differentiation using the β_ST_ index (Weir & Goudet, 2017) with the Hierfstat R package (Goudet, 1995), and measured the relationship between Hs, β_ST_, and the distance to the southernmost site using linear models. Hierfstat was also used to compute population’s Fis and compute 95% confidence intervals around the parameter using 1000 bootstraps over loci. Oceanographic distances between river mouths were measured with the marmap1.0.4 R package (Pante & Simon-Bouhet, 2013) to obtain more accurate distances compared to our previous analyses, where we used Euclidean distances between sample site coordinates.

We investigated population structure by performing a PCA on allele frequencies across loci with the program ade4 (Dray & Dufour 2007). A Multi-Dimensional Scaling plot based on estimates of pairwise relationships (identity-by-state) was constructed using Plink1.9 (Purcell et al., 2007) and plotted with the ggplot2 R package (Wickham, 2009). To better reveal fine scale population structure that may be spread across many PCA axes, we used variational autoencoders (VAEs) designed for population genetic inference (Battey, Coffing, & Kern, 2021). VAEs are unsupervised machine learning models that use a neural network (here, feed forward networks are used for the encoder and decoder parts) to regenerate the data and allow visualizing genetic data in a latent space that is a lower dimensional space than PCA while preserving geometry, unlike other recent methods designed for large datasets (Battey et al., 2021).

### Candidate Loci: Genotype Environment Association analyses (GEA)

We evaluated the extent to which genetic variation was explained by environmental variables using two approaches. First, we used a redundancy analysis (RDA) implemented in the vegan2.5.6 R package (Oksanen et al., 2019). The Russian Reka Saichik River population was excluded from this analysis due to a lack of climatic data and pronounced geographic discontinuity from North American populations. The Mad River population was also excluded due to uncertainty about its original provenance as well as samples from Bonneville Hatchery in oregon. We applied a minor allele count (MAC) of 15 (reason in supplementary materials), resulting in 59,115 SNPs with less than 5% missing data across 7,719 individuals. This MAC threshold was used given that outlier detection tests are not designed to accommodate rare variants. The following model was tested: allele frequency ∼ Normalized_Distance + Temperature + Precipitation + Elevation + Rock + Condition(Latitude). Here, Latitude was used as a conditioning variable in a partial RDA to control for the confounding effect of isolation by distance (Battey et al., 2020; Meirmans, 2012). Latitude was included as a way to partially control for population structure without “over-correcting” for this effect given that the pattern in our data arises due to both population structure and continuous latitudinal IBD following the southern expansions (Rougemont et al. 2020). Making a full correction for population structure may increase Type II error, as demonstrated by Lotterhos et al. (2015) and Forester et al. (2018) with simulations showing that correcting for population structure may result in the exclusion of some true signals of local adaptation and decrease the rate of true positives.

Model significance was tested using an ANOVA with 1000 permutations of the genetic data. A cut-off of three standard deviations from the mean loading was used to identify outlier SNPs on the significant RDA axes.Secondly, we applied the Genotype-Environment Association method based on latent factors mixed models (LFMM) representing residual population structure. This method is implemented in the R package LFMM1.5 (Caye, Jumentier, Lepeule, & François, 2019). We included Latitude as an additional explanatory variable in our model and excluded all outliers associated with this variable. A correction for multiple testing was performed following (Benjamini & Hochberg, 1995), and a given SNP was considered an outlier if its corrected p-value was below 0.01. Finally, we obtained functional annotations of outlier SNPs shared from the RDA and LFMM analyses using SnpEff v3.4 (Cingolani et al., 2012).

### Allele frequency distribution & parallelism

Given the importance of normalized distance as a candidate selective factor in our GEA analyses (see Results), we aimed to dissect the signal explained by adaptation to normalized migration distance. We thus compared populations performing «long» versus «short» distance migrations. Long-distance migration was defined by the distribution of normalized distances observed in our data (i.e., distance greater than 422km or elevation above 400 m, corresponding to the 15% upper quantile), whereas distance or elevation below this quantile corresponded to short migration. We then performed a PCA on outliers that were shared between RDA and LFMM to test the discriminatory power of these candidate SNPs between long *versus* short normalized distances.We worked only on shared outliers, as a way to further decrease the rate of false positive detection, without overly stringent corrections. If adaptation to migration distance shows parallel patterns at the genetic level, populations of long versus short migration distances should form distinct clusters, regardless of geography, and the same alleles at outlier loci should display parallel allele frequency shifts. Conversely, if geographic proximity plays a stronger role and/or if populations are adapting through allele frequency changes at different loci (non-parallel adaptation), then they should cluster by geographic groups. To further quantify whether adaptation to long-distance migration was parallel or non-parallel among geographic groups, we tested the following linear model: PC1 ∼ Normalized_Distance + Latitude + Normalized_Distance * Latitude. Here PC1 corresponds to the first (and only significant) axis of the previous PCA based on outliers and separating our populations. Under this model, if the Latitude term is significant and explains a large proportion of the variance, it would indicate non-parallel adaptation for migration distance among populations from different geographic areas. In contrast, if only the Normalized Distance term is significant, then this would suggest that populations are adapting through parallel changes at the molecular level, and independently of their geographic proximity. The interaction term represents both parallel and independent adaptation at the molecular level. We performed this analysis using all outliers associated with Normalized Distance detected by both RDA and LFMM and compared the results to a random subset of 400 SNPs, close to the number of SNPs identified by each method. We used the function sample in R after excluding all putative outliers to generate the random subset.

### Candidate loci: Differentiation based analysis

To further identify candidate loci involved in parallel (or non-parallel) adaptation to migration distance but displaying large allele frequency changes, we performed a differentiation-based genome scan taking advantage of multiple population pairs with long versus short normalized distances. Unlike GEA, which can detect subtle allele frequency changes and identify multiple covarying SNPs, differentiation-based genome scans have been developed to identify outlier SNPs that display large allele frequency changes (Lotterhos & Whitlock, 2015). Instead of relying on traditional genome scan methods that are affected by complex demography, we looked at patterns of shared outliers among multiple replicate pairs of populations sharing similar ancestry. To do so, we used pairs of populations from the same watershed with one population exhibiting long distance migration and one population exhibiting short distance migration. However, most upstream samples were not independent from each other, often flowing into the same downstream river, so that keeping all pairs would lead to pseudo-replication. This implies a strong reduction in the number of ‘approximately’ independent pairwise comparisons, reducing our sample set to four population pairs (detailed in **Table S03**). Two pairs were from the Alaska genetic group, one pair from the BC group, and one pair from the Thompson River. Analyses were performed using the Population Branch Statistic values (*PBS*) (Yi et al., 2010) in ANGSD (Korneliussen, Albrechtsen, & Nielsen, 2014). We identified outliers as those above the 0.90^th^ quantile of the *PBS* distribution (Rougemont et al., 2017). The number of outliers shared among population pairs was then computed to evaluate the extent of parallel adaptation to long-distance migration. If outliers are shared among several pairs of populations across the range, they are unlikely to be confounded by drift, providing stronger evidence for parallel adaptation.

### Accounting for recombination rate variation

Variation in recombination rate along the genome is known to influence the detection of outliers (Perrier & Charmantier, 2019). To test if the density of RDA, LFMM, and differentiation-based outliers increased in areas of low recombination, we estimated population-scaled estimates of recombination rate using LDhat (ρ = 4\**N*_*e*_*r where *N*_*e*_ represents the effective population size and r the recombination rate in cM/bp, **Supplementary Materials 1**). These estimates were obtained from 71 individuals sampled from 14 populations distributed from California to Alaska that were sequenced at the whole genome level (Rougemont et al., unpublished). We then tested for preferential enrichment of “hotspots” or “coldspots” of recombination in outliers (**Supplementary Materials 1**). We first tested if recombination hotspots (coldspots) contained more (less) outliers than “normally” recombining regions using χ^2^ tests. Linear mixed-effects models were then used to test the relationship between recombination rate and the distribution of outliers. The response variable was the SNP state, which was considered binomial (0 = non outlier, 1 = outlier) and the explanatory variable was the recombination rate. Chromosome identity was included as a random effect. Models were performed using the LME4 R package (Bates, Mächler, Bolker, & Walker, 2015). Our results revealed a significant association between detected outliers and recombination. Similarly, differentiation is known to be negatively correlated with recombination, so that many outliers in low recombination regions could be false positives (Booker et al., 2020; Perrier & Charmantier, 2019). The rate of recombination is proportional to the extent of LD across the chromosome (Ohta & Kimura, 1969): when recombination is small, high LD is expected whereas low LD is expected in regions of high recombination. Therefore, to control for the confounding effect of variation in recombination on differentiation-based outlier detection, we used an approach previously developed to account for large scale variation in LD (Perrier, Rougemont, & Charmantier, 2020). To do so, our previous population branch statistics (PBS) values computed across 1 megabase (1Mb) were used to estimate large-scale differentiation driven by large scale recombination variation. We subtracted these 1Mb window values from the PBS values of outliers identified in 50 kilobase (kb) windows to obtain a ΔPBS and corrected for variation in recombination rate. A low ΔPBS is indicative of a low difference between the background (associated to recombination) and the local window, possibly indicating a false positive local window. A large ΔPBS is indicative of a large difference between the background and the local window, so that variation in recombination should not be responsible for driving the pattern of differentiation. We computed the number of outlying windows that remained considering different ΔPBS thresholds. Finally, we used the approach in Rougemont et al., (2020) to test for correlations between PBS values and recombination. We reduced the multiple PBS values using the first axis of a PCA and tested for a correlation between the first PCA-PBS axis and recombination using mixed linear models including the chromosome identity as a random effect.

## Results

### Genetic structure mirrors geography within North American Coho salmon

Global patterns of population genetic structure corroborated our previous results (Rougemont et al., 2020). A PCA performed on 105,362 SNPs among 7,829 individuals across the entire North American Coho salmon range revealed that samples that are geographically closer are genetically more similar than more distant samples (**Fig. 1A, B, Fig. S03**). The first PC axis was correlated with longitude (r = 0.30, p <0.0001), whereas the second axis was correlated with latitude (r = 0.81, p < 0.0001), reflecting the decay of genetic similarity as a function of geographic distance (r = 0.35, p<0.0001, **Fig. S04**). While isolation by distance dominates at broad spatial scales, subtle population structure was also observed, likely due to the species’ homing tendency. This fine-scale structure was revealed using the VAE, which separated nearly every river within any given regional group **(Fig 2A**). As one example, on Haida Gwaii, all rivers clustered separately except for individuals from four rivers that formed a mixed cluster, consistent with signals of admixture **(Fig 2B, Fig S05**). These results are likely a product of river-scale homing typical in salmonids (Quinn, 1984).

**Figure 2:**
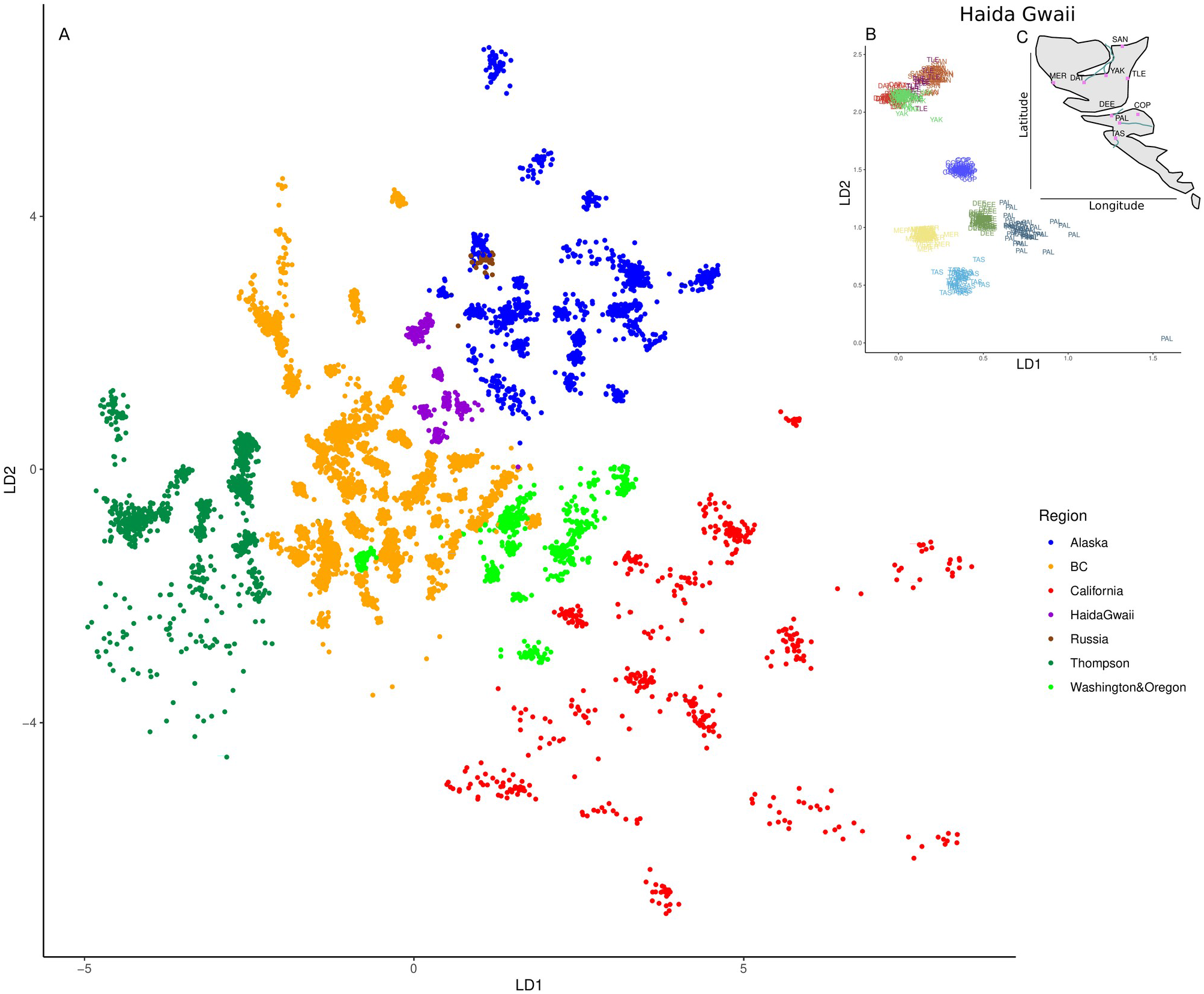
Variational Auto Encoders (VAE) revealing fine scale population structure at the river-scale level. Each point corresponds to an individual. A) Global structure inferred over the whole dataset; each color defines a major region; B) Zoom into one region (Haida Gwaii), highlighting closely related samples within river basins as well as fine scale structure; each colour defines a river on Haida Gwaii island with the three letters corresponding to the rvier abbreviation. C) Map of Haida Gwaii sample locality. Letters match those of river in panel b. Closer relatiosnhip in the geographical space is reflected in the location of the sample in the VAE. Each additional Region is highlighted separately in supplementary figure Fig S03.

### S*outhern populations are ancestral and the most genetically diverse*

Global patterns of variation in intra-population genetic diversity also corroborated our previous results (Rougemont et al., 2020). We observed a linear decrease in genetic diversity (Hs) as a function of the distance from the southernmost site (R^2^ = 0.39, p<0.0001, **Fig. S06A**). The Thompson and interior Alaskan populations deviated from the patterns observed for coastal populations (**Fig. S06A**), as expected, due to founding events during upstream postglacial recolonization. We therefore expected that this result would constitute a confounding factor in subsequent analyses (Fourcade et al., 2013).

Negative β_ST_ coefficients revealed that ancestral populations and those with high genetic diversity (**Fig. S06B**) were located in Cascadia and southern BC at the approximate southern boundary of the ice sheet during the last glaciation. A linear decrease in β_ST_ with distance from the south was observed (R^2^ = 0.39, *p* <0.0001, **Fig. S06B**). The Russian population did not display reduced genetic diversity or increased βst, as would be expected under a linear expansion from the southeastern Pacific coast. Removing the Russian population increased the R^2^ to 0.42 in our linear models.

### Genotype Environment Associations

The PCA on temperature-associated variables revealed four significant axes of variation that were kept for the RDA and LFMM analysis below (**Fig S02a**,**b**). Similarly the PCA on precipitation-associated variables (**Fig S02c**,**d**) revealed three significant axes enabling a decorrelation of the variables among them. The first four axes of the temperature PCA and first three axes of the precipitation PCA respectively captured 87% and 89% of the total variability. We verified that these axes as well as latitude, geology and normalized distance did not display strong correlation using the VIF and a correlation plot (**Fig S03**). Latitude displayed the highest VIF values (10.35), indicating significant co-variation with all other variables considered in our analyses (**Fig 3A**,**B**, **Fig S07**). All other variables displayed a VIF below 10 (**Table S04**) and were retained resulting in a total of 8 variables.

**Figure 3:**
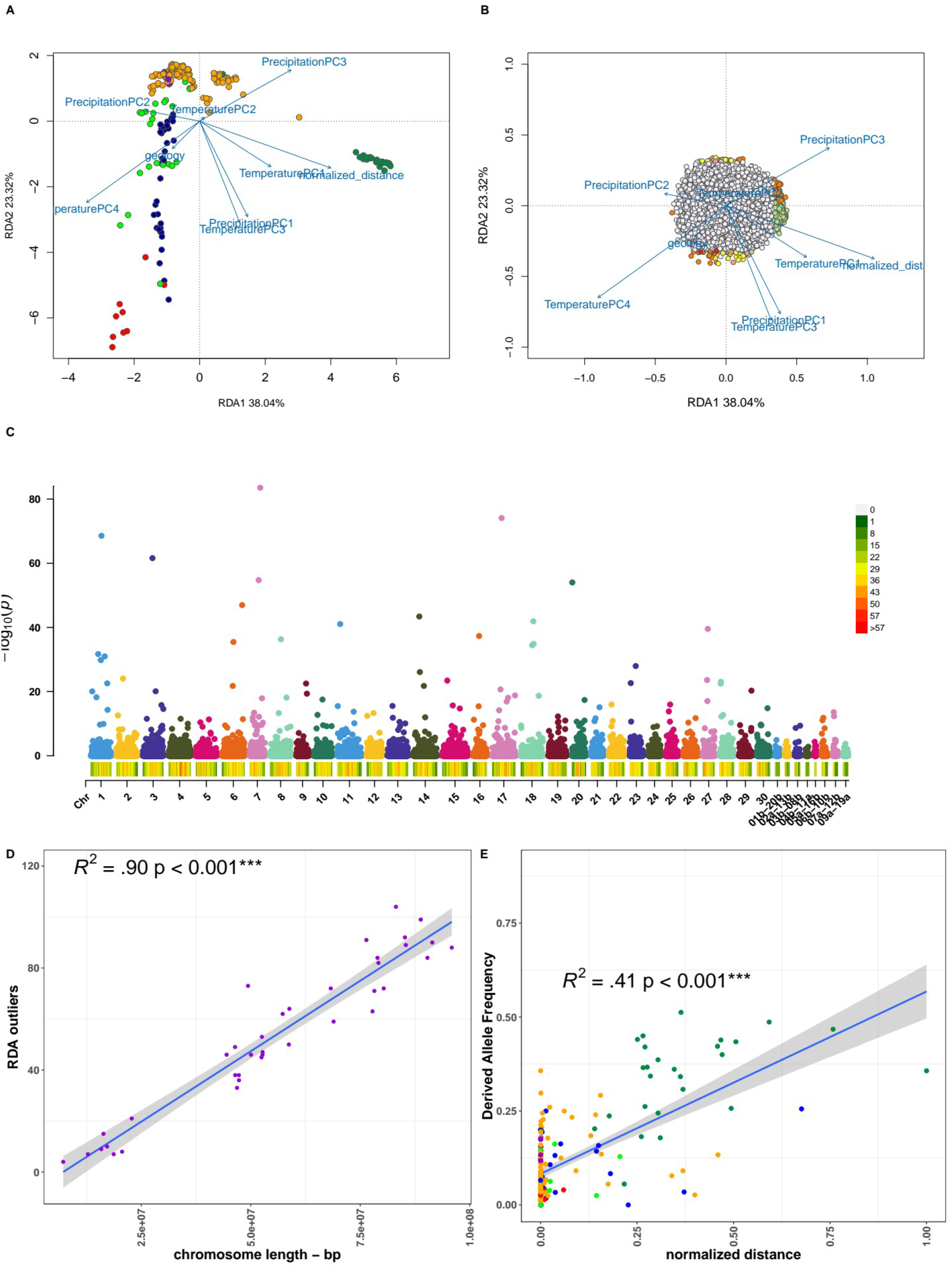
Landscape genomics results. A) RDA results showing the discrimination of populations along with environmental variables; points are populations colored according to the population from which they were sampled, B) RDA results showing outlier SNPs (colored points) associated with a given environmental variable. Environmental variable (Temperature 1, Temperature 2, Temperature 3 and Precipitation1, Precipitation2, Precipitation3) represents the first three PC axis of Temperature and Precipitation variable extracted from the worldClim database and decorellated through the PCA. The Grey dots correspond to non-outlier SNPs. C) Manhattan plots of -log10(p-value) for SNPs associated with migratory distance and identified by LFMM for K = 20 groups. X axis, left : Chr1 to Chr 30 : diploid chromosomes. X axis, right: Chr01b_20b to chr09a_19a: chromosomes with residual tetraploidy. SNPs falling on unassembled scaffolds were removed from the plot (n = 10 SNPs). D) Relationship between outlier count on each chromosome (y-axis) identified by RDA and chromosome length (x-axis). The shaded region displays 95% confidence intervals. E) Relationship between normalized distance (x-axis) and derived allele frequency (DAF) (y-axis) for the SNP in the vicinity of *epas1*. Each point represents a population and is colored according to each region of sampling.

The RDA permutation and ordinations tests revealed that all tested environmental and geographic variables were significant **(Table S04)**. The full model was significant and the environmental variables captured 11.18% of the total genotypic variance (DF = 2, p < 0.0001). The first seven RDA axes were significant, cumulatively capturing over 94% of the total variance. Outlier SNPs were identified on all significant axes (**Table S04**) with a total of 1,960 outlier SNPs being detected. Fifty-six percent (n = 1,105) of those outlier SNPs were associated with normalized distance to the spawning site and mostly associated with the Thompson River (**Fig 3A, Fig S08**). A further 521 and 292 SNPs were associated with temperature and precipitation, respectively, while 42 SNPs were associated with geology but mostly associated with other regions (**Fig 3A)**. The distribution of outlier SNPs across the genome was strongly correlated with chromosome length (adjusted R^2^ = 0.91, p <0.0001, **Fig 3D**).

Since strong postglacial founder effects may cause high rates of false positives, we also used a mixed latent-factor analysis in LFMM to identify shared outliers between different methods (Capblancq, Luu, Blum, & Bazin, 2018). As for RDA, a high number of outliers were associated with normalised migration distance (n = 1,465 SNPs or 65% of all detected outliers), and 352, 187, and 0 outliers were associated with precipitation, temperature and geology, respectively. Accordingly, 29% of outliers associated with normalized distance displayed strong selection signals (i.e. -log10(pvalues)> 5, **Fig 3C**), whereas other variables displayed a weaker signal (**Fig S09**). As with the RDA outliers, the outlier SNPs identified by LFMM were distributed throughout the genome with no strong peak. A total of 273 outliers were shared between the RDA and LFMM, among which 181 were associated with normalized distance, representing ∼62% of the total number of shared outliers. Of the remaining shared outliers, 12%(n = 36 were associated exclusively with precipitation, ∼9% (n=122) with temperature, and 18% (n=31) were associated,with different environmental variables (often Distance and another variable) by either RDA or LFMM, suggesting that these outliers may covary and jointly contribute to local adaptation in Coho salmon. Only one shared outlier was associated with geology.

### Parallelism in outlier distribution

The predominant role of normalized migration distance as a potentially strong selective factor based on the GEA results was investigated further using a PCA to test if populations were effectively separated into groups with long vs short distance migration. We performed a test on all shared outlier SNPs associated with this variable and shared between RDA and LFMM. To capture the strongest signal possible we restricted our RDA SNPs to those showing the strongest correlation with normalized distance (r > 0.65, pvalLFMM <0.001, n = 64 SNPs). As expected, only, the first PCA axis was significant and captured ∼83% of the total variance. This axis discriminated the Thompson River populations from all others, due to the fact that this river system comprises only “long normalized migration distance” populations (**Fig 4A**).A linear model revealed a highly significant effect of normalized distance (p < 2e^-16^) and of the interaction between latitude and distance (p < 2e^-16^), but no effect of latitude alone (p = 0.366), indicating that this confounding factor was well accounted for (**Table S05**). In comparison, a linear model performed on a random set of non-outlier loci revealed a very strong effect of geography (p < 2e^-16^) as well as normalized migration distance (p < 1e^-814^) and their interaction (p < .5.21e^-11^), highlighting the strong geographical component of our dataset that needed to be accounted for **(Fig S10)**. To identify additional outliers associated with geographic regions, we repeated the RDA and LFMM analyses excluding Thompson River populations. Levels of outliers sharing was of 237 shared SNPs between RDA and LFMM and 82 outlier SNPs were associated with normalized migration distance (**Table S06**). Of the, 82 SNPs associated with normalized distance that were shared between the two GEA analyses, 20 SNPs displayed a strong correlation (r > 0.5) with the significant RDA axes (**Fig S11** and **Table S06-S07**). This analysis clearly separated Alaskan populations displaying long normalized migration distance (e.g., Porcupine River, Mile Slough) from the remaining samples **(Fig S12)** and revealed outliers putatively under weaker selection that may have been missed when considering the whole dataset. Repeating the linear model approach described above on the set of 48 SNPs displaying the strongest correlations with normalized distance revealed no significant effect of latitude (p = 0.675) but a significant effect of normalized migration distance (p < 3e^-16^) and of the interaction between latitude and migration distance (p < 2e^-16^, R^2^ = 0.85, **Table S09**).

**Figure 4:**
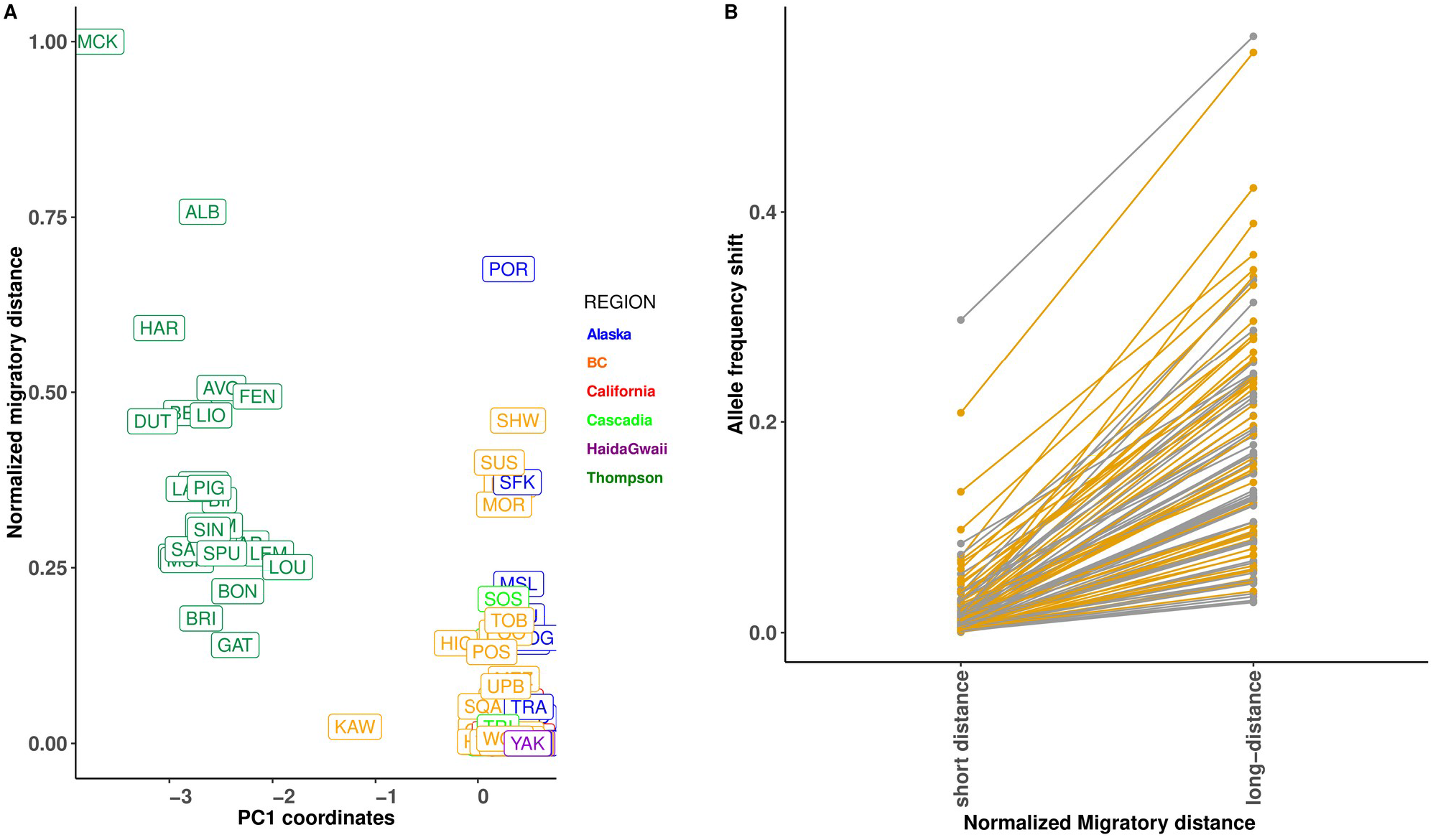
Outliers separate samples with short versus long normalized distance from Thompson and display strong parallel frequency changes across many SNPs. Left panel (A): PC1 coordinates based on all sampling locations and the set of the strongest outlier SNPs (n=47; r2 > 0.62) associated with normalized distance that were shared by RDA and LFMM analyses. The PC1 separates populations sampled according to the levels of genetic divergence associated with migratory distance. Each box represents a sample location and is colored according to its regional area of sampling. Right panel (B) displays parallel allele frequency shift for the 47 SNPs with the strongest signal of selection (r^2^ > 0.62). The two colors (grey and yellow) are chosen randomly for ease of visualisation. Rivers were divided into two groups representing «short» versus «long» Normalized Migratory Distance to evaluate allele frequency changes. See Figure S12 for results of the same analyses excluding Thompson River samples.

Averaged allele frequency changes (ΔAF) between the 41 strongest outliers associated with long *versus* short normalized migration distance were modest (mean ΔAF full dataset = 0.145, mean ΔAF without Thompson = 0.0565), but always in the same direction in the whole dataset (**Fig 4B**), even after removing Thompson populations (**Fig S12**). This result was in contrast with patterns observed with random SNPs, where allele frequency changes were stochastic (**Fig S10, Fig S12**). This suggests that allele frequency changes associated with migration distance responded in a similar way across multiple loci (similar directionality) among Coho salmon populations from different geographic regions.

### Signature of differentiation along the genome

Next, we looked at population differentiation among six pairs of long *versus* short normalized migration distances using PBS values to search for potential 10kb outlier windows (**Fig S13**). The top 10% of windows from the empirical PBS distribution identified a total of 3762 windows in the first pair of Alaskan samples, 3756 in the second pair, 3677 windows in BC, 3568 in ThompsonA total of 109 outlier windows were shared among all pairwise comparisons. Shared differentiation among population pairs was much higher at the regional level, with a total of 2,206 and 775 shared windows between the two pairs in Alaska and between the pair in Thompson and BC, respectively.. Similar results were obtained based on *F*_ST_ values (**Fig S14**).

### Confounding of genome scan and GEA outliers by recombination

A potential pitfall in interpreting outliers as signals of selection is that increased genetic differentiation (*F*_ST_ or PBS) may also be driven by the confounding effect of background selection (Burri, 2017) or by recombination rate variation (Booker et al. 2020). The ΔPBS value that we used to account for this potential confounder revealed that of the 35 shared windows among all pairs, eight displayed a ΔPBS > 0.2, while only height window displayed a ΔPBS > 0.3, indicating that background differentiation across windows was high and no particular window stands out with a strong signal of differentiation. Accordingly, linear mixed models revealed a low but significant correlation between PC-PBS and recombination rate (p < *2e*^*-16*^, R^2^c = 0.0017, R^2^m = 0.0096), supporting the effect of linked selection on the extent of observed genetic differentiation (Rougemont et al. 2020). The broad background *F*_ST_ distribution along the genome and among populations was illustrative of the effect of demography and recombination (**Fig S14**).

Similarly, regions of low recombination may be enriched in GEA outliers (Lotterhos, 2019). Here, shared outliers detected by both RDA and LFMM also fell preferentially in cold spots of recombination (χ^2^ = 21.11; df = 2, p *=0.04e*^*5*^). Accordingly, outlier windows displayed significantly lower recombination rates than «neutral» windows (W = 7,040,186 p *=* 0.00216, **Fig S15**) and the GLM confirmed this observation (p = 0.005, χ^2^ = 7.85; R^2^m = 0.0094, R2c = 0.0095). Therefore, it is possible that linked selection or areas of low recombination along the genome, which can enhance the effect of genetic drift, are not accurately modelled by our GEA and that these regions are enriched in false positives. In salmonids, genomic regions of residual tetraploidy may significantly influence our results. Indeed, we found that outlier SNPs in regions of residual tetraploidy did not differ from non-outlier in terms of recombination (see supplementary results).

### Functional annotation of outliers

SNPeff annotations were extracted for all candidate outliers shared by the combined RDA/LFMM analyses with and without the Thompson River populations (n = 494 SNPs) and associated with all environmental variables. Among them, ∼24% and ∼41% fell within intergenic regions and in introns, respectively. Moreover, 7% were composed of missense variants (n=34), 4% of 5’ or 3’ UTR regions, and the remaining SNPs were synonymous mutations (details in **Table S10**). A number of putatively important genes were identified (full list provided in **Table S10**). For instance, 34 SNPs were associated with various zinc finger proteins, 27 were close to transcription factors, and a number of other genes identified were involved in lipoprotein synthesis (e.g., lmf2 (missense variant)), growth and metabolic processes (e.g., akap12 (missense variant), spermatogenesis and one missense mutation was involved in cardiomyocite development (*Tmem88*). Furthermore, two of our four outliers with the lowest p-values identified only by LFMM were associated with metabolism and growth (i.e., a fatty acid-synthase like gene, and a tetraspanin-16 gene), the latter of which is known to be involved in cell development, activation, and growth, and its role in controlling trygliceride excess (hypertriglyceridemia) has been documented in humans (Weissglas-Volkov et al., 2013).

Lastly, outlier annotation revealed one SNP in the close vicinity (10.485kb) of *Epas1*, a strong candidate transcription factor gene known to be involved in adaptation to altitude and with a major role in cardiac performance (Yi et al. 2010). This SNP was associated with long normalized migration distance in all methods and displayed an increased derived allele frequency in all Thompson populations (DAF ∼0.4) and in the most distant Alaskan samples from the Porcupine River (DAF = 0.26) compared to populations with short distance migration. DAF was correlated with normalized migration distance (linear model R^2^ = 0.41, p < 2e^-16^ **Fig 3E, Fig S16**) and not confounded by either Latitude or Longitude (p > 0.05). Pairwise *F*_ST_ between populations experiencing long vs short normalized migration distance at this specific SNP was in the range 0.40-0.60 **(Table S11)**. The SNP is located in an intergenic region between *Epas1* and a *Pcgf5* gene that is known to be important for maintaining the transcriptionally repressive state of many genes throughout development (Yao et al., 2018). On this same chromosome, another downstream outlier gene (*agpt2*) is known to play an important role in oxygen performance (see Discussion).

## Discussion

This study is one of the largest landscape genomics efforts performed to date on a non-model species. It was made possible by a large multilateral sampling effort combined with an improved and well-annotated genome. The major objective was to document patterns of adaptive genomic variation across the species range and to identify the key environmental factors contributing to such adaptive variation. Below, we discuss these different issues and propose avenues of future research to expand beyond the current limitations of our study.

### Genetic variation mirrors geography and reveal fine scale homing

We confirmed the geographical component associated with the linear postglacial expansion from a southern refugium toward the north (Rougemont et al., 2020). The PCA axes were correlated with geography, and genetic relatedness between pairs of individuals decayed with geographic distance. These results indicate that genetic variation mirrors geography for Coho salmon, similar to human populations (Menozzi, Piazza, & Cavalli-Sforza, 1978; Novembre et al., 2008). As previously observed, some Californian samples displayed high genetic diversity while others displayed reduced diversity, strongly suggesting recent population declines in some of these rivers (Rougemont et al. 2020; Gilbert-Horvarth et al. 2016; Wilian et al. 2011). Accordingly, highest levels of genetic diversity and lower Bst were observed in Cascadia, rather than California, as already noted (Rougemont et al. 2021). We previously hypothesized that recent admixture between populations from several refugia resulted in an increased in genetic diversity at contact zone, rather than in the refugia themselves (Petit et al. 2001; Rougemont et al. 2020). Interestingly, our sample from a Russian population departs from a strictly linear decrease in diversity as one moves away from the South. This suggests either intercontinental gene flow following the colonization of this area in Russia, or that a second southern refugium existed in the western Pacific, which remains to be tested with the analysis of additional sampling in Russia. A last hypothesis is that this sample site may have been wrongly labelled as originating from Russia. In the absence of further data from western Pacific, none of these hypotheses can be excluded.

Beyond these broad-scale patterns, the analysis based on variational auto-encoders captured fine-scale structure that was not revealed in the PCA. Indeed, the PCA mainly enabled a genealogical interpretation of the results (McVean, 2009), here corresponding broadly to the divergence of the interior Thompson River populations from the rest of the dataset and to large scale isolation by distance from North to South. In contrast, variational auto-encoders revealed a biologically meaningful characteristic of Coho salmon. This approach clearly identified sampled populations that were more closely related to each other and may be useful for biomonitoring. In contrast to a PCA, for which the interpretation is well connected with the theory, it is less straightforward to draw theoretical expectations from variational auto-encoder, just as with most machine learning-based methods (but see discussion in Battey et al., 2020). Yet, we were able to detect biologically relevant patterns summarized in just two dimensions, highlighting the power of this approach. Similar to our previous work, we observed a decrease in genetic diversity and an increase in population differentiation toward the North, yielding further support for the «out-of-Cascadia» scenario followed by postglacial founder events moving northward (Rougemont et al., 2020).

### Multiple environmental variables potentially driving local adaptation

We sought to identify the major environmental variables responsible for driving local adaptation in Coho salmon throughout their North American range. Normalized migratory distance, interpreted as a selective environment that would result in variation in migratory phenotypes, is frequently included in GEA studies of salmonids. Distance is expected to have major selective effects, given the energy expenditure and physiological requirements involved in reproductive migration (Bernatchez & Dodson, 2011; Eliason et al., 2011). Indeed, our GEA suggested a major role of normalized migratory distance (representing 73% of all outliers), a variable commonly identified as a driver of local adaptation in other salmonid species that undergo variable anadromous migratory distances. This is the case in Chinook salmon, *O. tshawytscha* (Brieuc, Ono, Drinan, & Naish, 2015; Hecht et al., 2015); Steelhead trout (Hess, Zendt, Matala, & Narum, 2016; Micheletti et al., 2018) and Arctic char, *Salvelinus alpinus* (Moore et al. 2017). A smaller yet non-negligible role of precipitation (10% of all outliers) and temperature (7%) was also observed, while geology appears to play a relatively minor role, in contrast with other studies in salmonids (Quéméré et al., 2016). The role of precipitation and temperature as selective forces have also been demonstrated in other salmonids (Bourret et al., 2013; Dallaire et al., 2021; Eliason et al., 2011; Micheletti et al., 2018) as well as in other taxa, including in plants (Leroy et al., 2020) and *Drosophila* (Kapun, Fabian, Goudet, & Flatt, 2016). In salmonids, Michelleti et al. (2018) suggested that extremely high precipitation is likely to impose an additional energetic cost during migration to spawning grounds. Conversely, low flow combined with warm temperature may impede access to spawning grounds and constitute a particularly strong selective force (Gilbert & Tierney, 2018). Temperature, combined with migratory distance, is likely to be a selective force along the distribution range of Coho salmon. For instance, Alaskan populations face warm temperatures during the upstream migration (17.9°C), sometimes combined with long migratory distance (e.g. Porcupine River), which we suggest, may jointly impose strong selective pressures potentially driving local adaptation (Olsen et al., 2011). Similarly, southern populations in California experience strong selective pressures for local adaptation due to warm temperatures, which, if combined with dry periods, may select for resistance to these factors that have important implications for the adaptation of these populations to climate change. Our sampling design may have limited power to detect loci associated with long distance migration across the whole dataset. Indeed, the majority of our sampling localities associated with long distance migration occurs in the Thompson, five locations were in Alaska whereas only one river with long distance migration was located in the southern range Additional sampling focussing on this particular question would therefore be relevant. Another limit of our current study is its reliance on RADseq data, which represent a limited portion of the genome, with approximately 1 SNP every ∼40kbp here. Given the extent of LD, it is plausible that we have missed relevant signals of small to intermediate effect as well (Lowry et al. 2017). Further whole genome sequencing on a set of carefully chosen populations from our study would provide additional clues regarding the genetic architecture of adaptation to long distance migration.

### Recombination impacts on genome scans

The distribution of GEA outliers was correlated with chromosome length, which was recently proposed as an indication for a polygenic basis of adaptation (Salmón et al., 2021). However, this result may also be related to the lower rates of recombination in larger chromosomes and may indicate an increased rate of false positives due to linked selection. Indeed, Rougemont et al., (2020) previously showed that recombination rates were associated with chromosome length in Coho salmon (R^2^ = 0.48, p <0.0001) and that the distribution of deleterious variants was correlated with chromosome length, suggesting linked selection. We have shown that nucleotide diversity is correlated with recombination rate (Rougemont et al., 2020), a pattern that may also be a signal of linked selection (Burri, 2017). *F*_*ST*_-based outlier tests are known to be influenced by variation in recombination rate since the neutral estimates of differentiation will be overly inflated in regions of low genetic diversity (Cruickshank & Hahn, 2014). In contrast, our GEAs consider subtle allele frequency changes related to environmental variation. In comparison to *F*_*ST*_-based outlier tests, recombination rate variation is not expected to significantly bias the observed genotype-environment associations (see also Lotterhos 2019). Yet, our finding that a greater number of GEA outliers are typically observed in regions of lower recombination suggests that false positive detections may occur in these regions. Yet, our finding that a greater number of GEA (LFMM only) outliers are typically observed in regions of lower recombination suggests that false positive detections may occur in these regions. In addition, since RADseq outliers are more likely to be linked SNPs rather than causal variants, this increase in outliers in low recombination regions could also be due to greater retained linkage between RAD SNPs and causal variants. More theoretical work should be undertaken to help interpret these results and derive expectations regarding rates of false positives in GEAs. In addition accounting for the duplicated nature of salmonids genome and contrasted rates of recombination in duplicated and non-duplicated region would require further invsetigations (Allendorf et al., 2015; Brieuc, Waters, Seeb, & Naish, 2014; Danzmann et al., 2008; Kodama, Brieuc, Devlin, Hard, & Naish, 2014).

Similarly, for the *F*_*ST*_-based analyses, we found a positive correlation between *F*_*ST*_/PBS and recombination rate. This result is indicative of pervasive linked selection (Burri, 2017) or extended regions of low recombination potentially resulting in an excess of false positives (Booker et al., 2020). We attempted to reduce these potential biases using a LD-based approach by correcting PBS values according to values observed across large windows (Perrier et al., 2020). In this way, windows showing a large differentiation relative to the background are more likely to represent true positive detections. This strongly reduced our set of candidate windows. Therefore, it appears that low recombination rate is likely generating false positives that are difficult to account for. A potential way forward would be to perform differentiation-based scans by considering discrete classes of recombination, but the limited number of markers available from RADseq in most studies using this genotyping technique prevents such an approach. We suggest that linked selection in areas of low recombination is likely to shape the landscape of divergence among Coho salmon populations. Similar results have been observed in other salmonids (Lehnert, Kess, Bentzen, Clément, & Bradbury, 2020; Rougemont & Bernatchez, 2018) and other species (Burri et al. 2017, Perrier et al, 2020). A potential way to refine *Fst-*based analyses would be through neutral envelope estimation obtained by combining coalescent and spatially explicit forward simulations (Haller & Messer, 2019; Kelleher, Etheridge, & McVean, 2016) with heterogeneous recombination. Considering the above, we did not interpret further our *F*_*ST*_-based results and only focused on our GEA-based results, which should be less affected by recombination rate variation (Lotterhos 2019).

### Partial parallelism and subtle allele frequency change

As expected under the hypothesis of polygenic selective effects, our set of shared SNPs across GEA methods displayed modest and parallel allele frequency changes among all populations (**Fig 4b**). This suggests an important role of standing genetic variation enabling repeated allele frequency shifts at similar loci (Höllinger, Pennings, & Hermisson, 2019). An important caveat, shared by many studies relying on RADseq to investigate parallelism (e.g. LeMoan et al. 2016; Rougemont et al. 2017; Jacobs et al. 2020) is that candidate outliers are more likely to be SNPs linked to causal variants, rather than causal variants themselves (Lowry et al. 2017). In this case, true parallel patterns for causal loci may be masked if outliers represent linked neutral variation that varies among geographically structured populations. With this caveat in mind, we also observed that parallelism was strong at the regional scale (e.g., within Alaska n = 2 or within the Thompson and BC (n=2)) and decreased with distance. This is expected, as geographically distant populations showed increased levels of genetic divergence (Bohutínská et al., 2021). For example, a recent study of local adaptation by flowering time in *Arabidopsis* compared global and regional subsamples and reported that genetic effects are distinct in different locally adapted populations (Lopez-Arboleda, Reinert, Nordborg, & Korte, 2021). We detected one outlier, a missense variant (*tmem88* on chr9), involved in cardiomyocite development that was present at appreciable frequency only in Porcupine River (AF = 0.22). This population is the most extreme in terms of long migratory distance, coldest temperature, and lowest precipitation and illustrate the fact that if adaptation is really local then several outliers should only be population specific. However, the broad sampling we performed here is likely more powerful to detect allelic variants contributing to adaptation globally rather than locally. One caveat is that this population is also at the expansion front and many deleterious alleles may have surfed to high frequencies as previously observed (Rougemont et al., 2020, Rougemont et al. in prep). Therefore, it is also possible that neutral alleles have also surfed to high frequency. The functional role of these missense variants would need further investigation.

In our large dataset, we expected adaptation to occur through many genes of small effect (polygenic adaptation) rather than through a single gene of major effect as reported previously in Rainbow Trout (Hess et al., 2016; Pearse et al., 2019), Atlantic salmon (Ayllon et al., 2015; Barson et al., 2015), and Chinook salmon(Narum Shawn R. et al., 2018; Prince et al., 2017; Thompson et al., 2020). Moreover, in these species, a binary difference in phenotypes (e.g. late vs early maturing phenotype in Chinook or Rainbow trout) was present and the expression of these alternative phenotypes are associated with alternative alleles on single genes. However, such binary phenotypic differences in life history traits has never reported in Coho salmon (Quinn et al. 2016).None of our outlier SNPs mapped exactly onto genes of strong effect previously reported in several of these studies (e.g. *Rock1* and *Greb1L* associated with return timing, and *Vgll3* and *six6*, associated with age at maturity). This is similar to recent findings by (Waters et al., 2021), who did not identify any significant associations between the latter two genes and age at maturity in Coho salmon. Moreover, our study did not focus on variation in life history and phenotypic traits among populations, and therefore we did not investigate similar molecular targets of selection.

Among the different genes with a putative role, the transcription factor Endothelial Pas Domain Protein 1 gene (*Epas1)* deserves further investigation. The importance of *Epas1* for altitude adaptation in Tibetan humans (Yi et al., 2010), dogs (Wang et al., 2014), mice (Schweizer et al., 2019) and birds (Xiong et al., 2020) have been well established, with evidence for spatially varying selection. More specifically, this gene is involved in improving cardiac function, muscle activity, oxygen availability, and aerobic-anaerobic metabolism under hypoxia (Henderson et al., 2005). Interestingly, physiological differences in cardiac responses between locally adapted populations was demonstrated in a congeneric species, Sockeye salmon (*O. nerka*) (Eliason et al., 2011) as well as in other Pacific salmon including *O. mykiss and O. kisutch* (Eliason & Farrell, 2016). In particular, coastal populations with short upriver migration were shown to display a low aerobic and cardiac scope when compared to populations with a long and challenging spawning migration (Eliason, Wilson, Farrell, Cooke, & Hinch, 2013). Moreover, salmon facing more challenging spawning migration conditions displayed greater aerobic scope, larger heart, and better coronary supply resulting in adaptation at very local scale (Eliason et al., 2011). Therefore, in fishes, challenges to migration imposed by the environment are very likely to select for individuals displaying higher cardiovascular and muscular abilities, resulting in populations that are locally adapted to reach distant and high-elevation spawning sites (Eliason et al., 2011, 2013). Here, we identify for the first time strong candidate loci potentially underlying the genetic basis of these adaptations.

## Conclusion

Our results first confirm our previous inference of a south to north, as well as west to east, gradient of genetic diversity and population divergence. This pattern reflects historical contingency associated with a single refugium followed by postglacial expansion. Our findings provide evidence for adaptation of Coho salmon to their local environment, especially with respect to migratory distance and elevation, and to a lesser extent temperature and precipitation PCA axes. We also observed parallel variation in allele frequencies at a small number of outlier loci associated with environmental variation in migratory pathways, but the extent of parallelism decreased with increasing distance among populations, suggesting that local adaptation to similar environmental pressures involves different genomic regions in different populations. Finally, in contrast to previous studies on salmonids that identified a few genes of major effect associated with sexual maturity or migration timing, we failed to detect a significant role for these genes in Coho, but rather found more support for polygenic adaptation, as also reported in other studies (Bernatchez, 2016). Since our study used a partial representation of the genome, it may have missed the signals reported at well-known genes. Therefore, future studies involving whole genome sequencing and functional validation are needed to further unravel the genetic basis of local adaptation in Coho salmon.

## Acknowledgments

We thank B. Bougas, A. Perrault-Payette, and C. Hernandez for their support in the laboratory. Computations were mainly performed on the Colosse (Calcul Quebec), as well as Graham and Cedar (Compute Canada) servers. This research was carried out in conjunction with EPIC4 (Enhanced Production in Coho: Culture, Community, Catch), a project supported in part by University Laval and the government of Canada through Genome Canada, Genome British Columbia, and Genome Quebec.

## Data availability

*All scripts to reproduce the analyses at any step can be fond on the first author’s github page (https://github.com/QuentinRougemont/coho_ldscp_genomics). RAW data have been deposited on NCBI under project PRJNA647050. Vcf files are available on dryad at* https://doi.org/10.5061/dryad.bzkh189b2.

