## Supplemental Methods and Figure for "Long distance migration is a major factor driving local adaptation at continental scale in Coho Salmon"

### Supplementary Methods

#### Molecular Laboratory protocol

Genomic DNA was extracted with a salt-extraction protocol modified from Aljanabi and Martinez (1997). Sample quality and concentration were checked on 1% agarose gels and a NanoDrop 2000 spectrophotometer (Thermo Scientific). DNA concentration was normalized to 20 ng/µl. Libraries were constructed following a double-digest RAD (restriction-site-associated DNA sequencing; Andrews, Good, Miller, Luikart, & Hohenlohe, 2016) protocol modified from Mascher, Wu, Amand, Stein, and Poland (2013). Genomic DNA was then digested with two enzymes (PstI and MspI) by incubating at 37°C for 2 hr followed by enzyme inactivation by incubation at 65°C for 20 min. A unique individual barcode was ligated to each sample using a ligation master mix including T4 ligase along with sequencing adaptors. The ligation reaction was completed at 22°C for 2 hr followed by 65°C for 20 min for enzyme inactivation. Samples were multiplexed (n = 48 individuals) to ensure fish from each sampling location were sequenced on a minimum of six different multiplexes to avoid pool effects. Libraries were size-selected using a BluePippin prep (Sage Science), amplified by PCR, and sequenced on an Ion Proton P1v2 chip.

#### Bioinformatic analyses

Each read was trimmed to 80 bp using cutadapt along with barcode removal (Martin, 2011). The "process_radtags" module of Stacks v2 was used for demultiplexing (Rochette et al. 2019). Reads were then aligned to the Coho salmon reference genome v2 (GCF_002021735.2) using bwa-mem 0.7.13 Li, 2013 and filtered with samtools v1.7 (Li et al. 2009) to keep reads with a mapping quality above 20, remove supplementary alignment and unmapped read. Variants were then called with Stacks v2. A mapq of 20 was required in gstacks and the module "populations" was run (requiring a given SNP to be present in at least 60% of individuals in each population) to produce a vcf file that was subsequently filtered stringently.

First, we excluded any SNP with a sequencing depth below 10 and above 120 based on the mean read depth computed over all individuals for each SNP (vcftools –site-mean-depth). The upper threshold on depth was used to control for PCR duplicate as well as putative paralogs. Although, in other studies we previously filtered our data based on excess heterozygosity to minimize paralogy and control for hardy-weinberg disequilibrium (e.g. Rougemont et al. 2019) we did not performed such filtering, because SNPs under strong selection in a single population may strongly depart from HWE and show unusual Fis pattern if not polymorphic in all other populations (e.g. this would be expected if SNP associated to long distance migration sweep to fixation in only a single population of 15 individuals out of the 7,829 individuals). We removed any individual with more than 20% of missing genotypes, reducing our dataset from 7,945 individuals to 7,829 individuals and removed any SNPs missing in more than 85% of the sample. Next, we kept a single SNP per locus as most of our analyses are not designed to account for linkage disequilibrium. Ultimately, we created two datasets (**Fig S1)**: 1) one for unbiased estimates of demographic parameters and 2) another for our GEA and associated analyses.

1. The first dataset was made of all **7,829** individuals and further stringently filtered to keep SNPs present in at least 95% of individuals. This vcf contained **105,362** SNPs with a mean sequencing depth of **28** (i.e. depth between 10 and 120 to control for paralogs and PCR duplicates) and a genotyping rate of **0.9876**.

2. The second dataset was designed for GEA, so we removed the samples from Russia as well as Bonneville (BNV), and kept SNPs with a minor allele count of at least 15, present in at least 95% of individuals and with a sequencing depth between 10 and 120 to control for paralogs and PCR duplicates.

This dataset comprises **7,759** individuals with **59,453 SNPs** a mean sequencing depth of **28.9** and a genotyping rate of **0.986**.

The reason to filter on minor allele count (MAC) rather than minor allele frequency (MAF) is as follows: Given the very large size of our dataset, a minor allele frequency of 1% would require a SNP to be polymorphic in at least 78 individuals, which is far more than the average sample size per river (n = 36). Given the high genomic structure of populations with long distance migration (e.g. Porcupine River in Yukon) any SNP fixed in this population would be removed from our dataset with a 1% MAF. Therefore, we seek to remove some low frequency variants that are unlikely to contribute to adaptation using a MAC of 15, as a trade-off between removing true biological signal and noise.

Finally, we created a third dataset by excluding the samples from the Thompson watershed due to their high divergence. Excluding these samples resulted in 189 populations and 56369 SNPs present in 6,758 individuals with a genotyping rate of 0.987, a MAC of 15, and a sequencing depth between 10 and 120 (mean = 28.84).

#### Analysis of Environmental data

Given the variability in temperature and precipitation across the study area, we included the mean, minimum, maximum, range and standard deviation of 19 climatic variables resulting in 95 variables associated with temperature (°C) and precipitation (mm).
These were extracted from the WordclimV2.0 database (Fick & Hijmans, 2017) for the period 1970-2000. We reduced these data to a set of uncorrelated variables using two separate principal components analysis (PCA) on the temperature and precipitation variables and retained the significant axes of variation (the first three axes in each case). The PCAs were performed using the R package ade4 (Dray & Dufour, 2007). The first four axes of the temperature PCA and three axes of the precipitation PCA respectively captured 86.9% and 88.6% of the total variability. We also extracted geological variables (rock type, geological era) from the USGS database (Garrity & Soller 2009) since geology has been identified as influencing population genetic structure in other salmonids (e.g., Bourret et al., 2013; Quéméré et al., 2016). The geological era variable was distributed unevenly across categories and was removed so that only the rock type variable (4 categories: Metamorphic, Plutonic, Sedimentary and Volcanic) was retained.
Coho from different populations throughout their range undertake a range of freshwater migratory distances (from a few kms to >2,300 km) to reach their breeding sites (**Table S1**), which we predict should result in differential selective pressures across populations. In addition to distance to the breeding site, elevation is expected to exert a strong selective pressure by increasing migration harshness (e.g., Bernatchez & Dodson 1987) with possible differences at the gene level when comparing low elevation to high elevation sites. Therefore, we computed the product of river length and altitude gain, standardized to a mean of zero and with a standard deviation of one (Moore et al. 2017), which will be referred to hereafter as «normalized distance». We verified the collinearity among all predictor variables prior to our analysis using the variance inflation factor (VIF). No predictor displayed a VIF greater than 10 so all were retained for GEA analyses.

#### *Inference of recombination rate variation with LDhat*

Whole genome SNP calling

To infer fine scale recombination rate, we used 71 individuals representing 14 populations from California to Alaska (Rougemont et al., 2020). Each individual was sequenced on an Illumina platform using paired-end 150 bp reads targeting a mean coverage of 30x. Reads were processed using fastp for trimming, bwa mem v0.7.13 (Li, Ruan, & Durbin, 2008) for mapping, samtools v1.7 (Li et al., 2009) requiring a minimum quality of 10, and picard to remove duplicates (http://broadinstitute.github.io/picard/). Then SNP calling was performed using GATK (McKenna et al., 2010). Genotypes were filtered for depth between 10 and 100 reads to remove low confidence genotypes including potential paralogs displaying high coverage. Then, following GATK Best Practices we excluded all sites that did not match the following criterion: MQ < 30, QD < 2, FS > 60, MQRankSum < -20, ReadPosRankSum < 10, ReadPosRankSum > 10. We also generated a vcf file using the samtools mpileup pipeline, merging individuals with bcftools and performing the same stringent filtering as with the vcf constructed using GATK. Finally, we also generated a separate vcf file using the emit-all-sites option to call variable and invariable sites across the whole genome. This vcf file was used in the sliding window analysis below to test the effect of linked selection. The whole pipeline is available on github (https://github.com/QuentinRougemont/gatk_haplotype).

Recombination rate estimate:

We used LDHat software (Auton & McVean, 2007) to estimate (population) effective recombination rates ρ = 4.Ne.r where r represents the recombination rate per generation and Ne is the effective population size) along the genome. Unphased genotypes were converted into LDHat format using vcftools with a minimum MAF of 10% since only common variants are useful for such inferences. Following the authors' guidelines, the genome was split into chunks of 2,000 SNPs with overlapping windows of 500 SNPs to compute recombination rate and data were then merged together. We measured recombination rates for each river as well as globally including all populations except the populations from the Thompson R. watershed, which were too divergent from the remaining samples. The pipeline to reproduce the analysis is available on github (https://github.com/QuentinRougemont/Ldhat_workflow).

Identification of recombination's coldspot:

In our analyses we seek to identify coldspot versus hotspots of recombination. To define a coldspot, we computed a lower bound that we defined as the mean population scaled recombination rate (ρ): ρ - 5*standard errors for each chromosome separately. Similarly, 'hotspots" of recombination were identified using an upper bound defined as mean ρ + 5*standard errors. We classified each 250 kb windows as a coldspot if the average recombination rate of a given window fell below the threshold (or above for hotspots).

Taking into account residual tetraploidy:

Genome from salmonids have undergone a whole genome duplication 80-100 million yeas ago (Macqueen & Johnston, 2014) and are ongoing rediploidization. Therefore, parts of the genome still exhibit residual tetraploidy. We tested whether these regions of the genome displayed a similar pattern of increased rates of outliers in low areas of recombination as compared to diploid regions.

To do so, we separated outliers according to whether they fall in region of residual tetraploidy or not and performed our mixed linear models (see methods in main text) for these regions separately. There was only 8 SNPs (out of 303 SNPs) present in region of residual tetraploidy when considering outliers shared between LFMM and our RDA. Therefore, to statistically test for an effect of residual tetraploidy we considered all outlier from LFMM (n = 67 SNPs on chromosomes with residual tetraploidy out of 2,257 outliers) and all outlier from our RDA separately (n = 81 SNPs on chromosomes with residual tetraploidy out of 2,054 outliers).

When considering LFMM results, there was no difference in recombination between outlier and non-outlier when considering areas of residual tetraploidy ($\rho_{non\text{-}outlier}$ = 0.536 , $\rho_{outlier}$ = 0.516, wilcoxon-test = 74,634 p-value = 0.09, glmer p-value = 0.32).

Similarly, when considering RDA results, there was no difference in recombination between outlier and non-outlier when considering areas of residual tetraploidy ($\rho_{non\text{-}outlier}$ = 0.535 , $\rho_{outlier}$ = 0.537, wilcoxon-test = 102,538 p-value = 0.69, anova p-value = 0.87).

Therefore, excluding chromosomes with residual tetraploidy, resulted in stronger differences in rates of recombination at outlier versus non-outlier SNPs compared to our main results (see main text). (e.g. RDA: $\rho_{non\text{-}outlier}$ = 0.721 , $\rho_{outlier}$ = 0.695, wilcoxon-test = 51,164,364, p-value = 0.0005; LFMM: $\rho_{non\text{-}outlier}$ = 0.721 , $\rho_{outlier}$ = 0.703, wilcoxon-test = 55,657,661, p-value = 0.009)

***Derived Allele Frequency (DAF) identification***

The procedure of Rougemont et al. (2020) was used to estimate derived alleles. To do so, we used the genomes of three outgroup species, (chinook salmon, rainbow trout and Atlantic salmon) to classify SNPs as ancestral or derived. Whole genome data for the chinook salmon (n = 3 individuals) were provided by B. Koop (unpublished), whereas rainbow trout (n = 5) and Atlantic salmon (n = 5) data were downloaded from NCBI Sequence Read Archive (rainbow trout, SRA, Bioproject: SRP117091; Salmo salar SRA Bioproject: SRP059652). Every individual was aligned against the Coho salmon V2 reference genome (GCF_002021745.2) using GATK HaplotypCaller and calling every SNP (invariant and variant) using the EMIT_ALL_SITES mode. We then constructed a custom Python script (available on github) and determined the ancestral state of the GBS SNPs if 1) the SNP was homozygous in at least two of the three outgroups, and 2) match one of the two alleles identified in Coho salmon. Otherwise, the site was inferred as missing and was not used in subsequent analyses.

**Supplementary Tables:**

**Table S01:** Dataset characteristics, associated summary statistics (Ho = Observed Heterozygostiy, Hs = Gene diversity, Bst = Differentiation index from Weir & Goudet, Fis = inbreeding coefficient) and environmental variables (normalized distance = distance * altitude, normalized to 1), geology = Rock type: Metamorphic, Plutonic, Sedimentary and Volcanic, BioClimatic variable (raw values) for the 19 bioclimatic variable available from the WorldClimV.2.0 database and reduced to three significants PCA axis for Precipitations and Temperature respectively.

**Table S01b:** Description of the 19 bioclimatic variable.

**Table S02:** Filtering steps to produce the different dataset used in our different analyses.

**Table S03:** List of populations chosen for pairwise Fst/PBS comparison of short vs long normalized distance across different geographic areas

**Table S04:** RDA Results including all sites. Showed is the significance of environmental variable and of each RDA axis.

**Table S05:** Linear models between outliers associated with the distance and environmental variables

**Table S06**: a) outliers shared between conditioned RDA and LFMM for the full dataset
B outliers shared between conditioned RDA and LFMM without the Thompson sites

**Table S07**: RDA Results without Thompson Sites. Showed is the significance of environmental variable and of each RDA axis.

**Table S08:** List of shared outliers between LFMM and RDA along with chromosome, position and SNP_ID information; pval_BH displays the pvalue after BH correction and LFMM_var indicates the associated variable detected by LFMM; axis indicates the RDA axis on which the SNP association was detected, the loading of each SNP as well as the correlation of each environmental variable with the SNP is displayed, along with the best predictor (i.e., strongest association with the outlier SNP) and its correlation.

**Table S09:** Linear models between outliers associated with the distance and environmental variables. The Thompson samples were excluded.

**Table S10:** Annotation of each outlier obtained with SNPeff.

**Table S11:** Fst at the SNP associated to EPAS1 when comparing long versus short distance population pairs.

### Supplementary Figures:

**Figure S01: Sampling site at the regional level.**

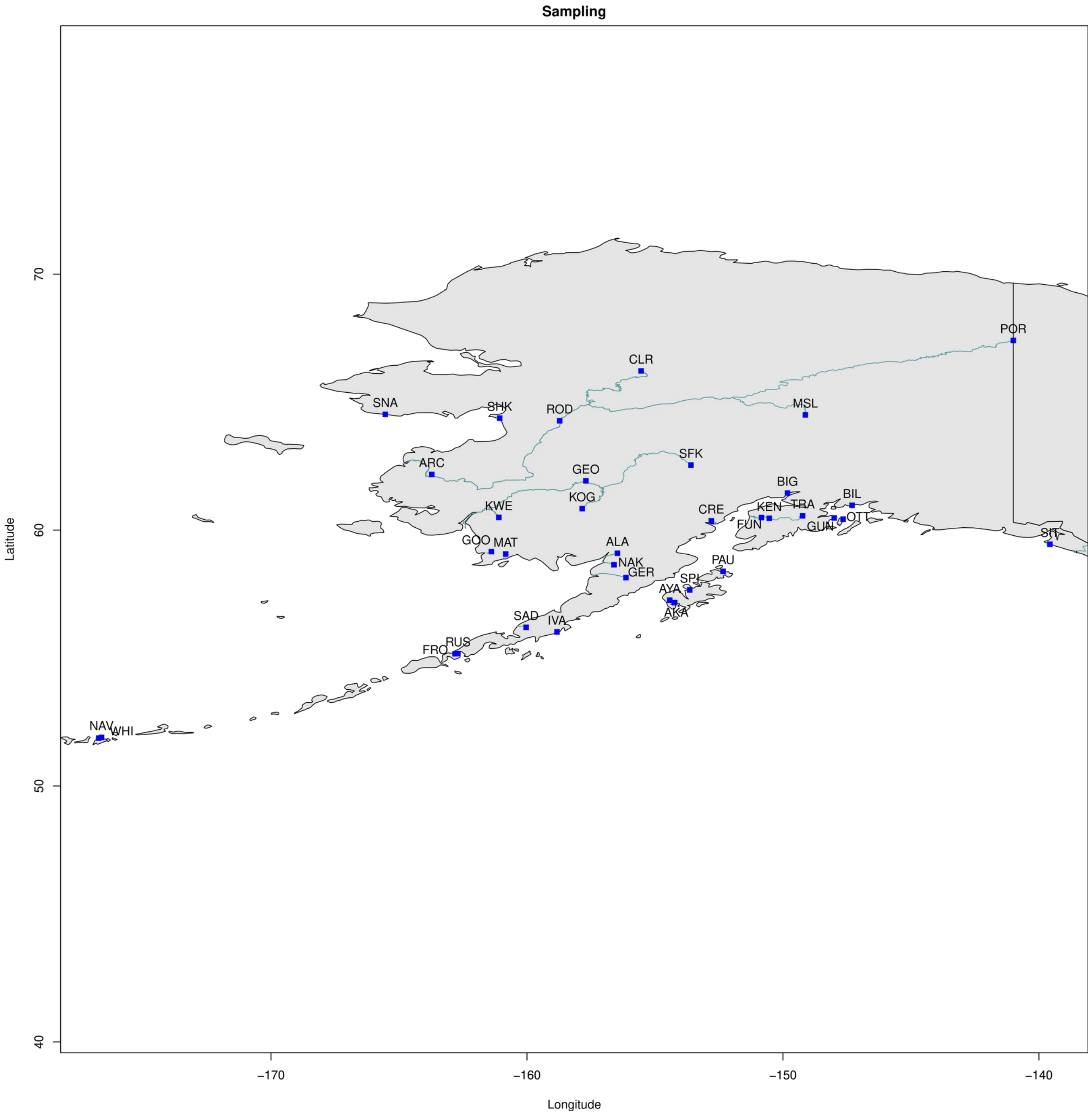


**Figure S01 a) details of the population sampling in Alaska. Abbreviation matches the code provided in table S01.**

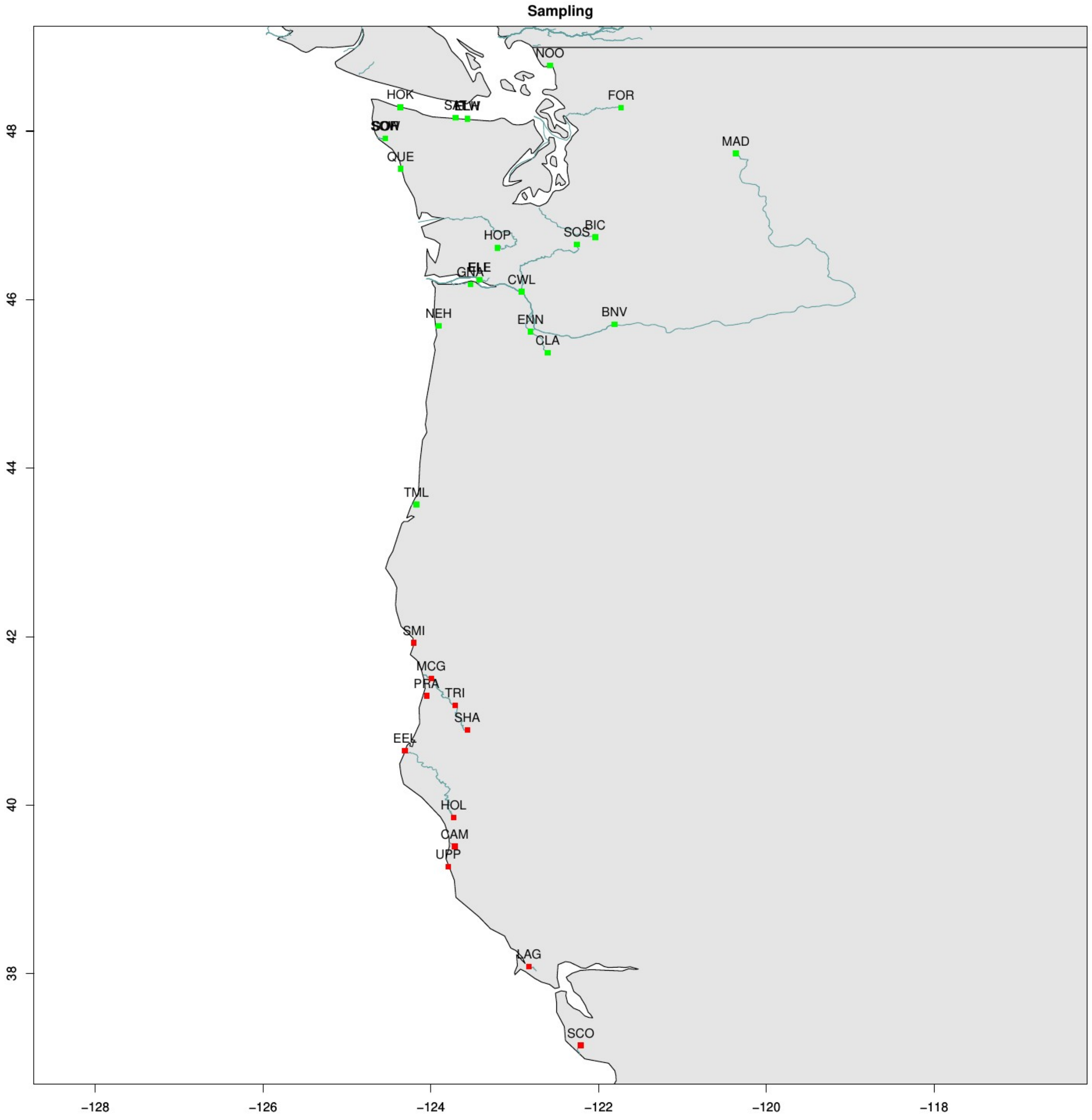


**Figure S01 b) details of the population sampling in California and Cascadia. Abbreviation matches the code provided in table S01.**

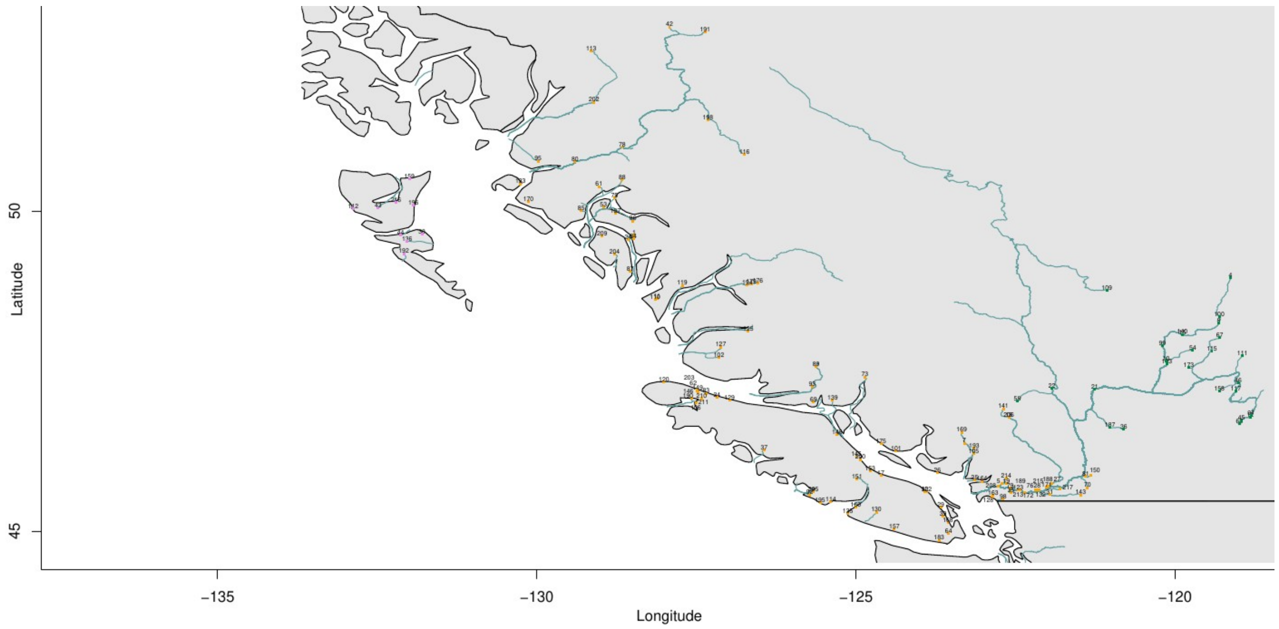


**Figure S01c) Details of the population sampling in BC including Thompson and Haida Gwaii. Number matches the code provided in table S01. A zoomable view of the map is provided on the Readme of github at [https://github.com/QuentinRougemont/coho_ldscp_genomics/](https://github.com/QuentinRougemont/coho_ldscp_genomics/)**

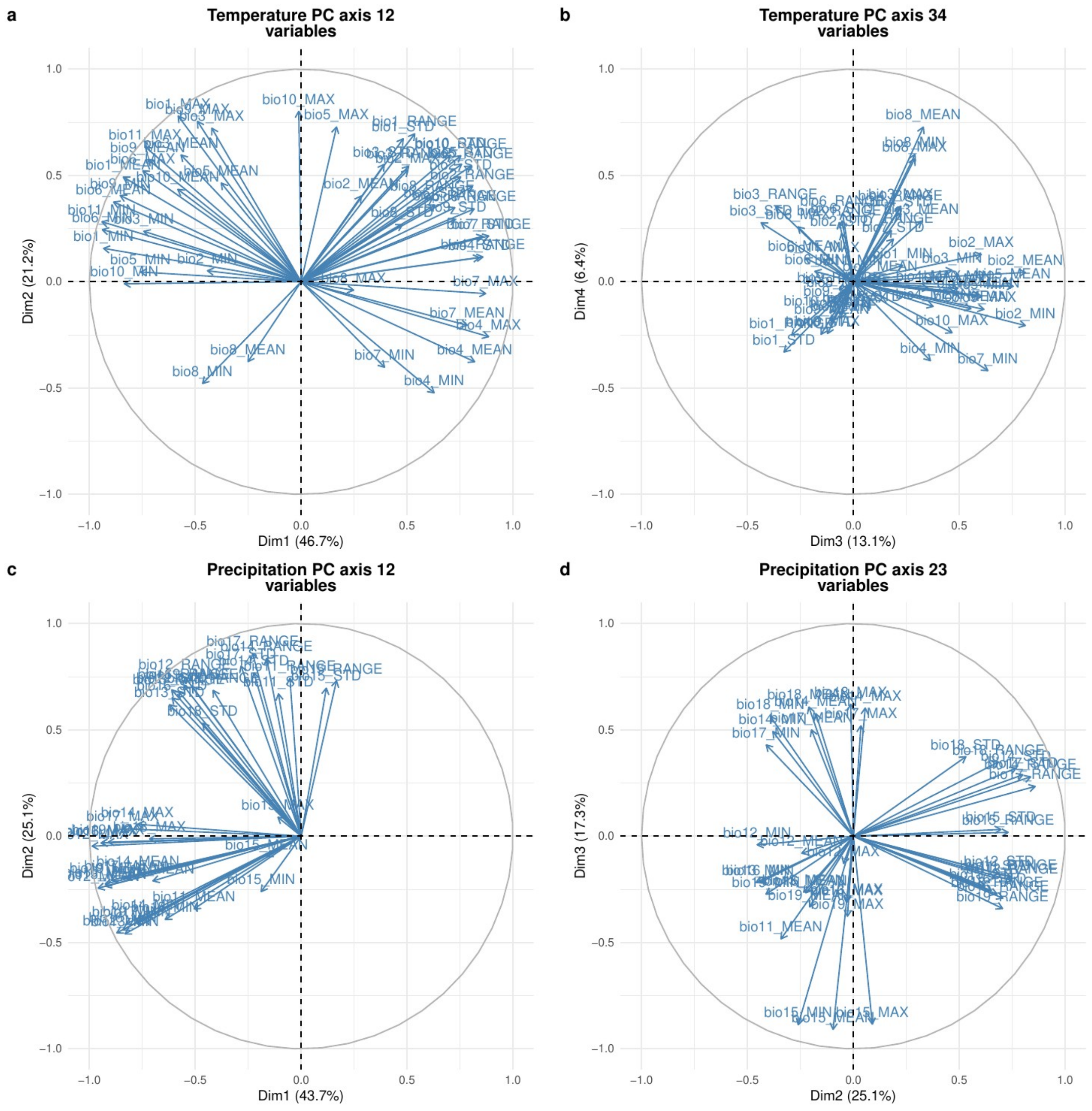


**Figure S02: PCA of environemental variables.** Only the significant axis are displyaed.

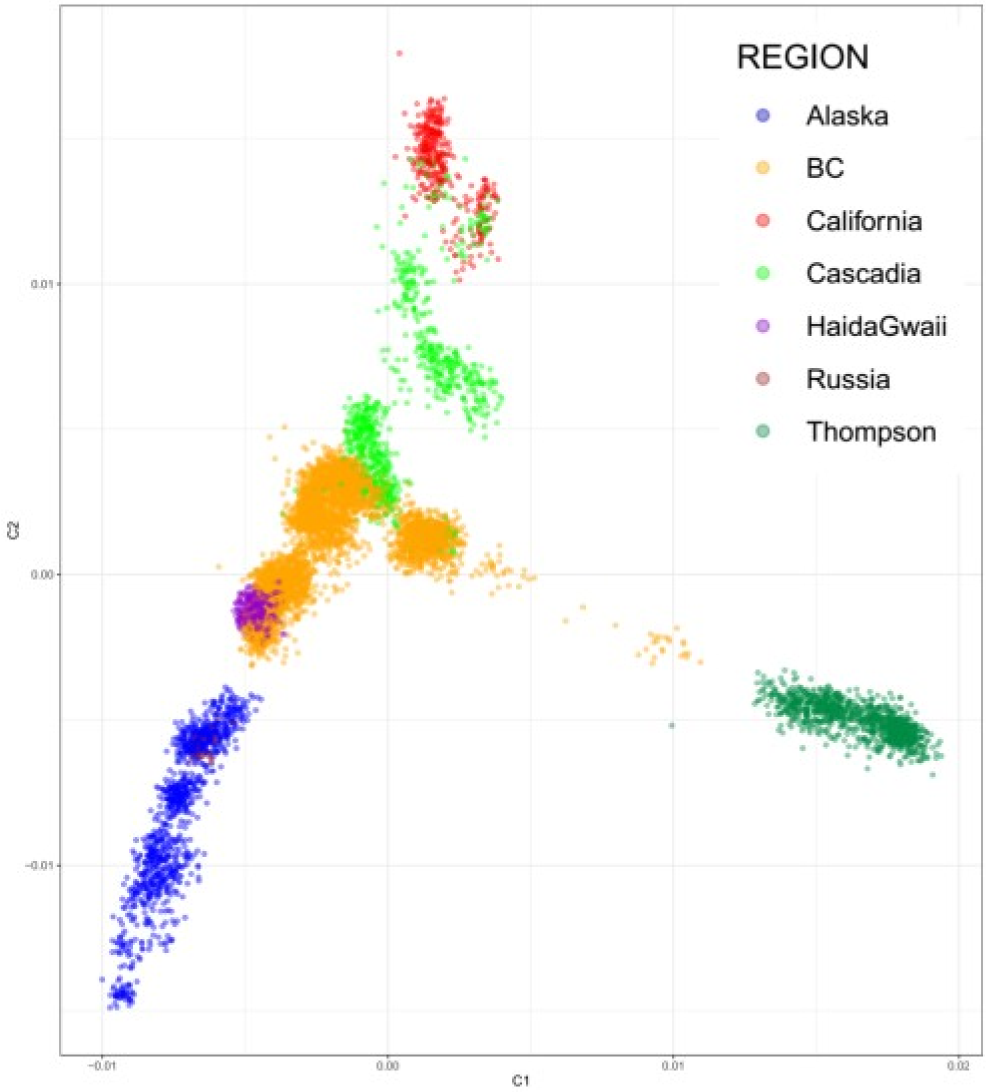


**Figure S03**: Plot of Identity by State for all individuals. Each point represents an individual sample and each individual is coloured by major regional group

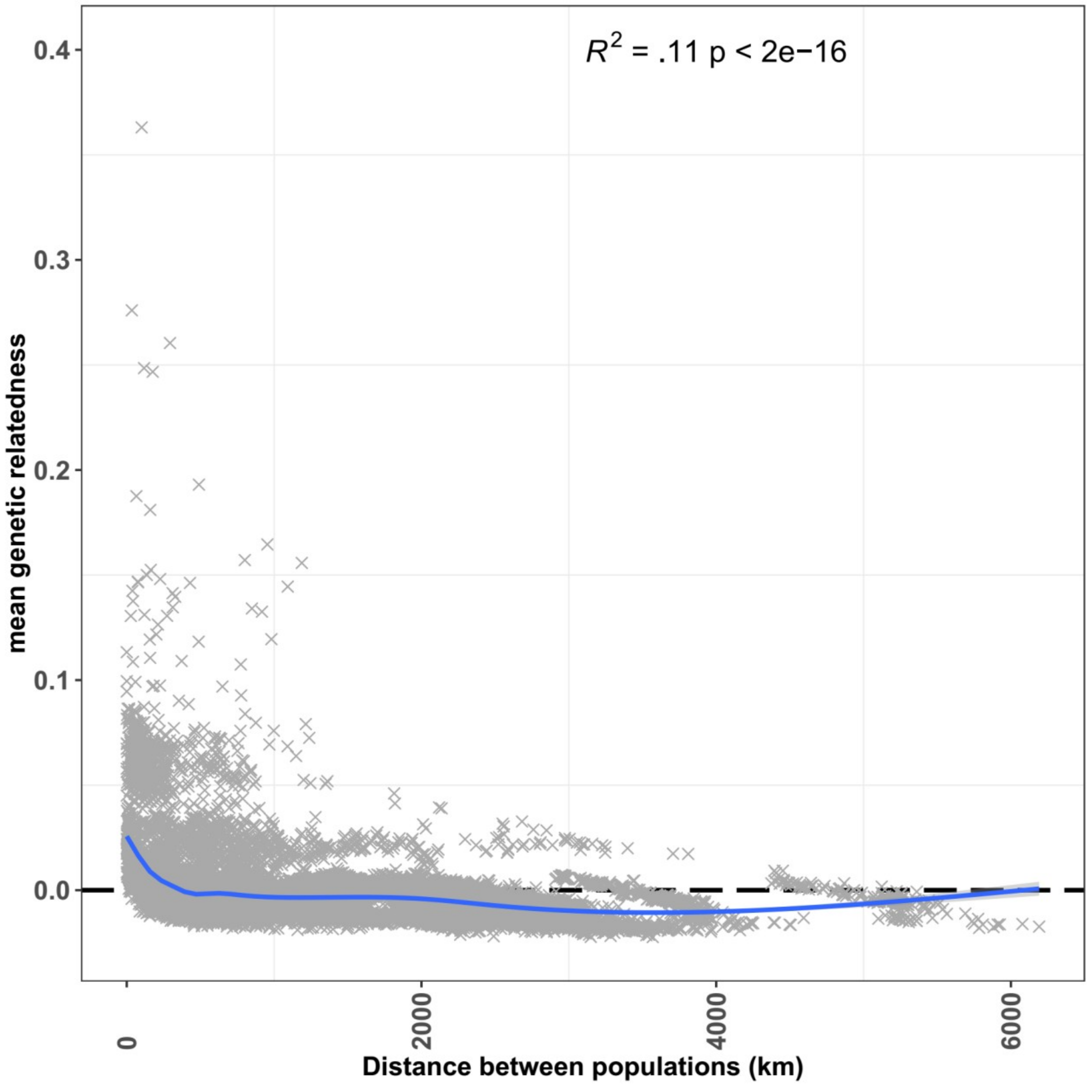


**Fig S04:** Significant correlation between relatdenness decay and distance between pairs of individuals.

**Fig S05 Variational Auto-Encoder revealed both fine scale structure and mixing among river in different region.** Results obtained after plotting each region separately. Region are plotted from South to North with A) Cascadia, B) California, C) BC, D) Thompson and E) Alaska. See Fig2 for HaidaGwaii and the global plot. Each individual is plotted by its river of sampling  abbreviated using a three digit code (see Table S1 for all correspondances) for ease of discrimination.
High quality figure are provided on github.

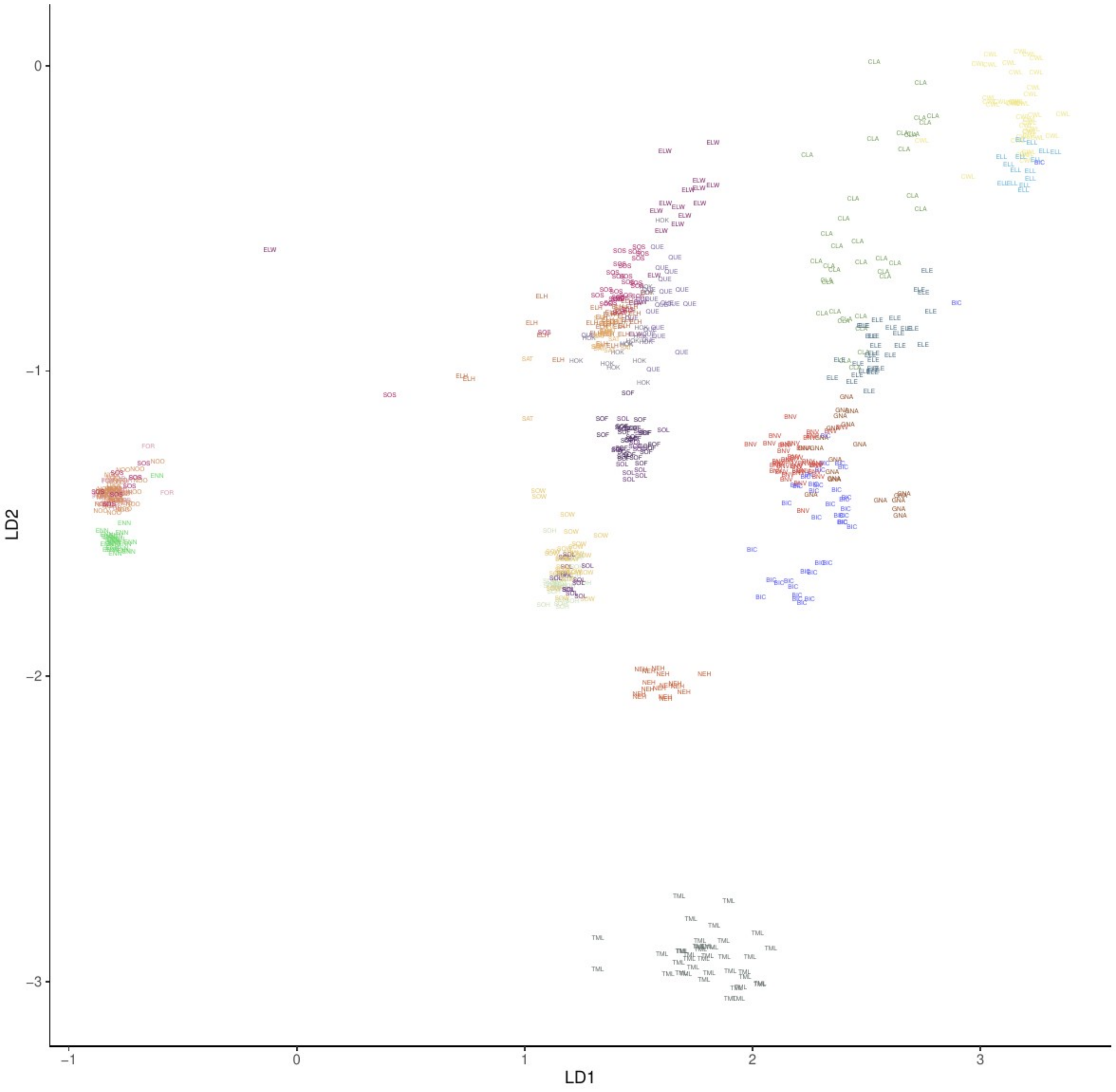


#### A) Cascadia

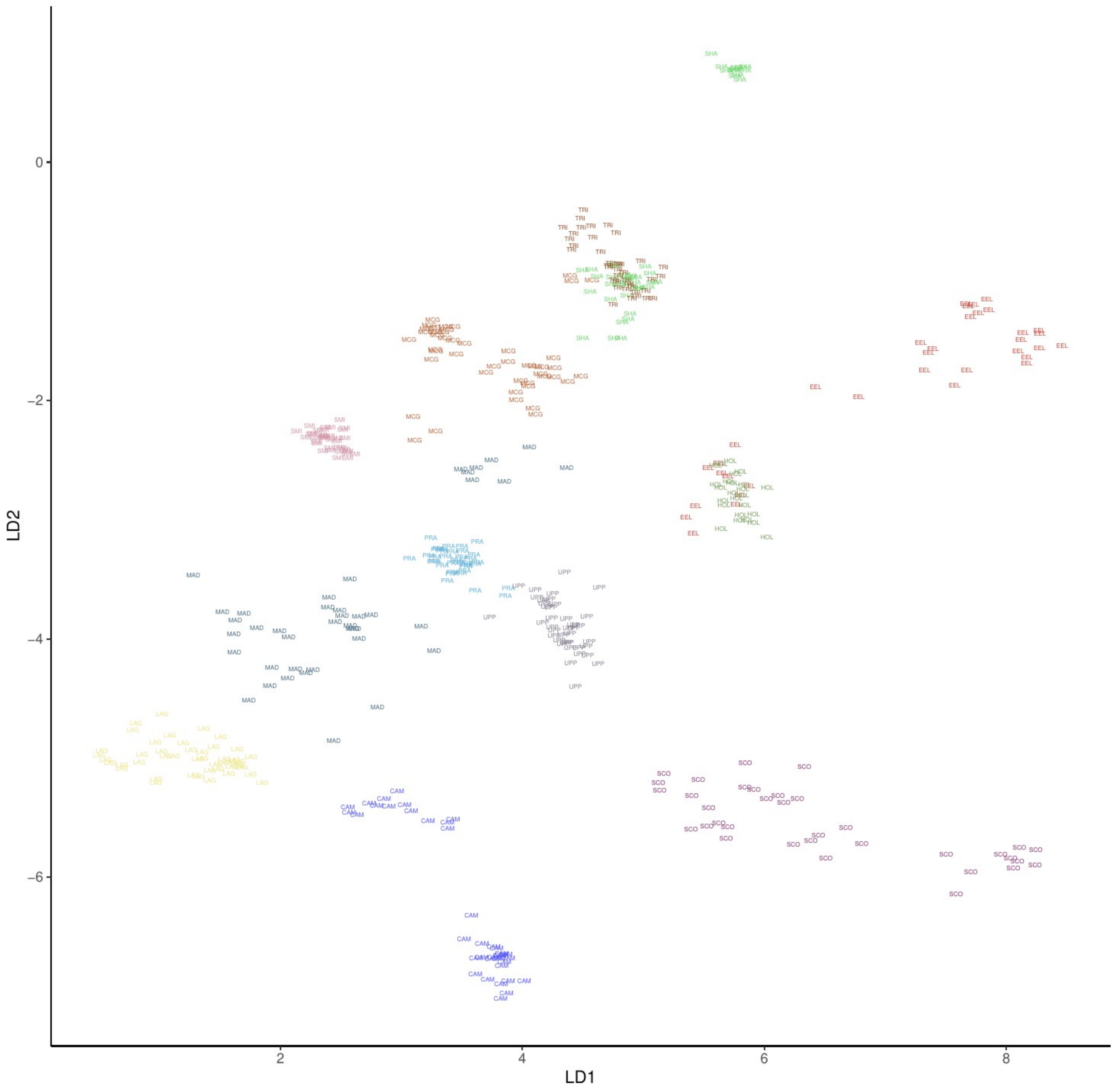


**B) California**

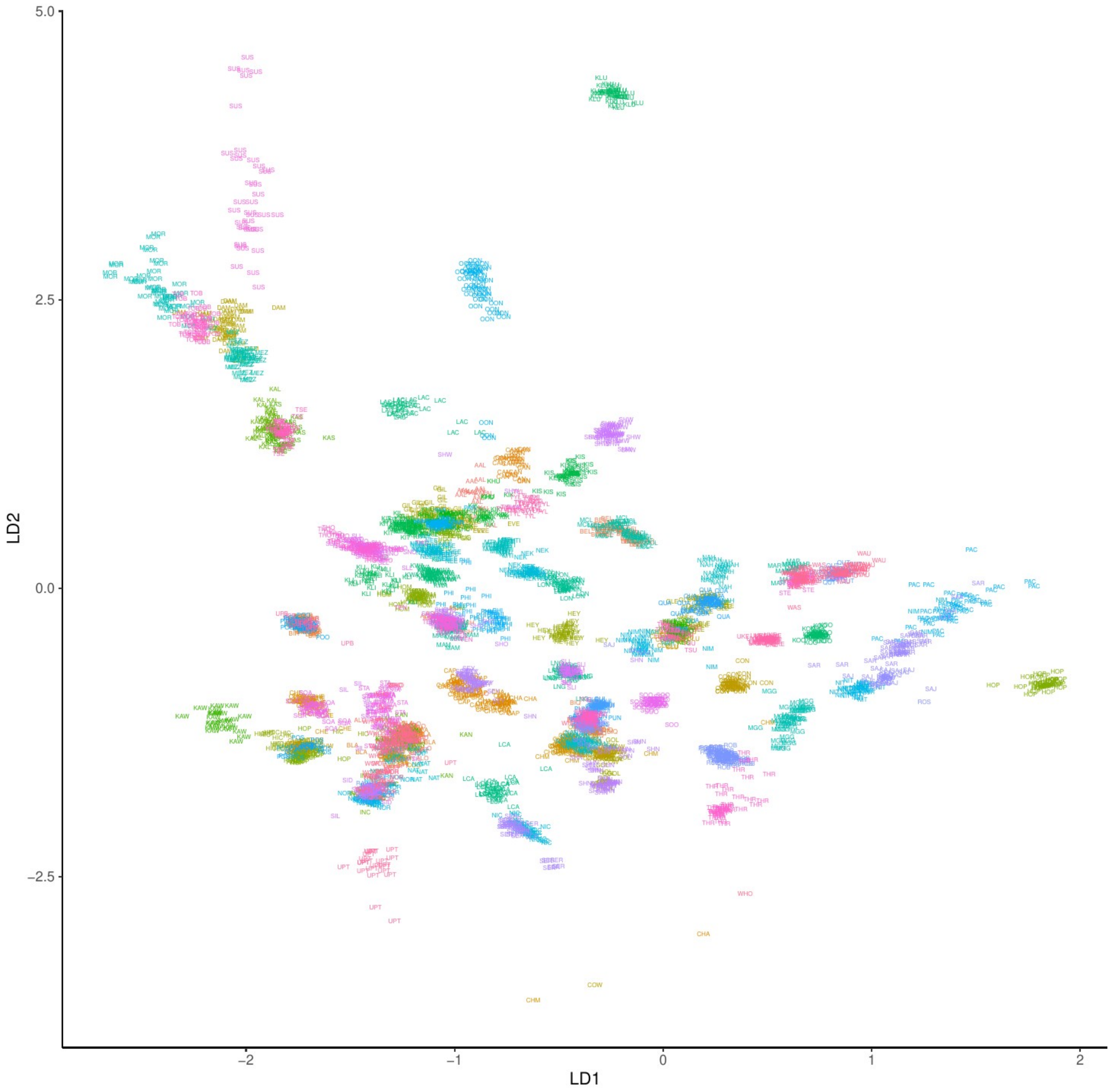


C) **BC**

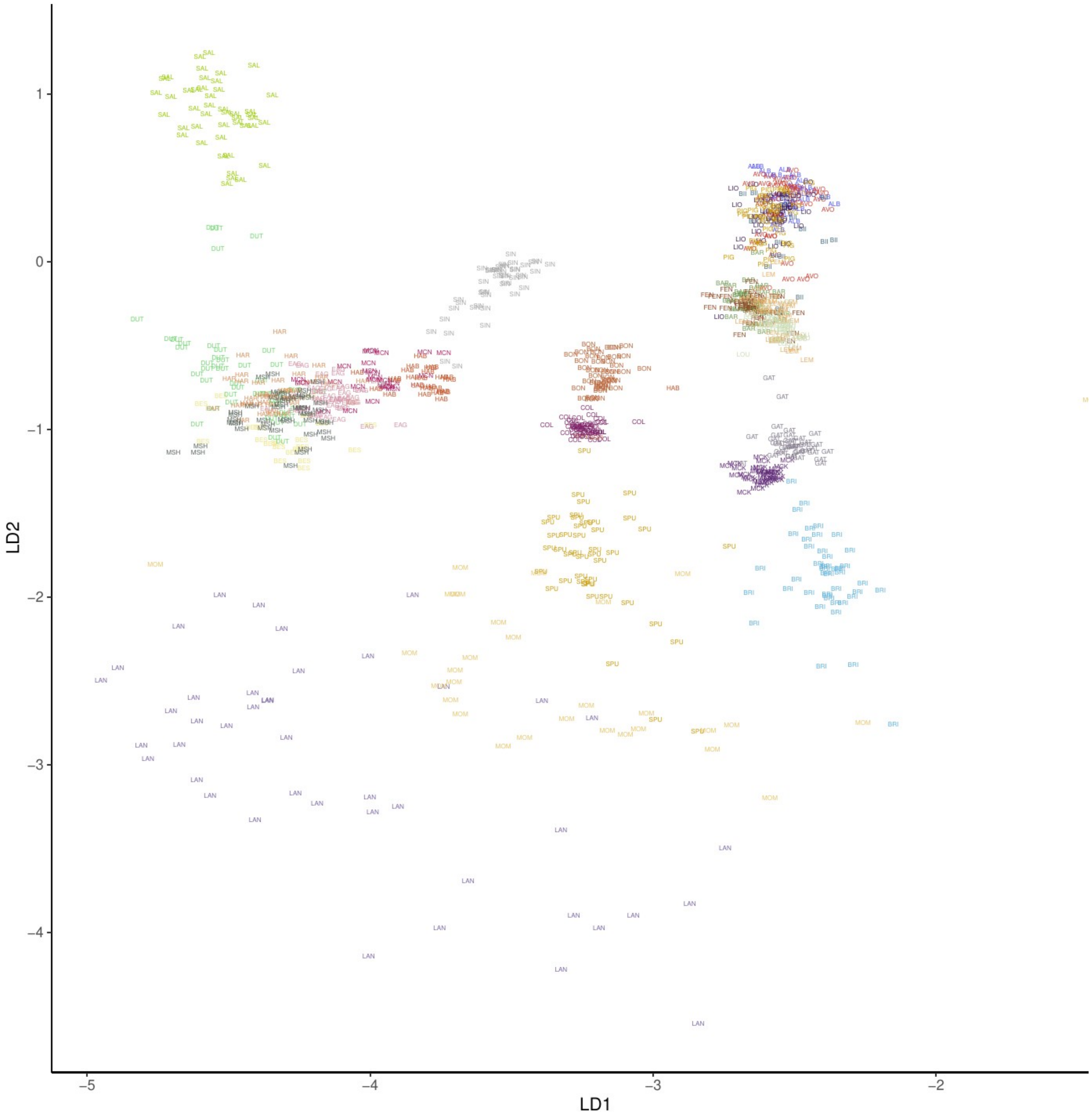


D) **Thompson**

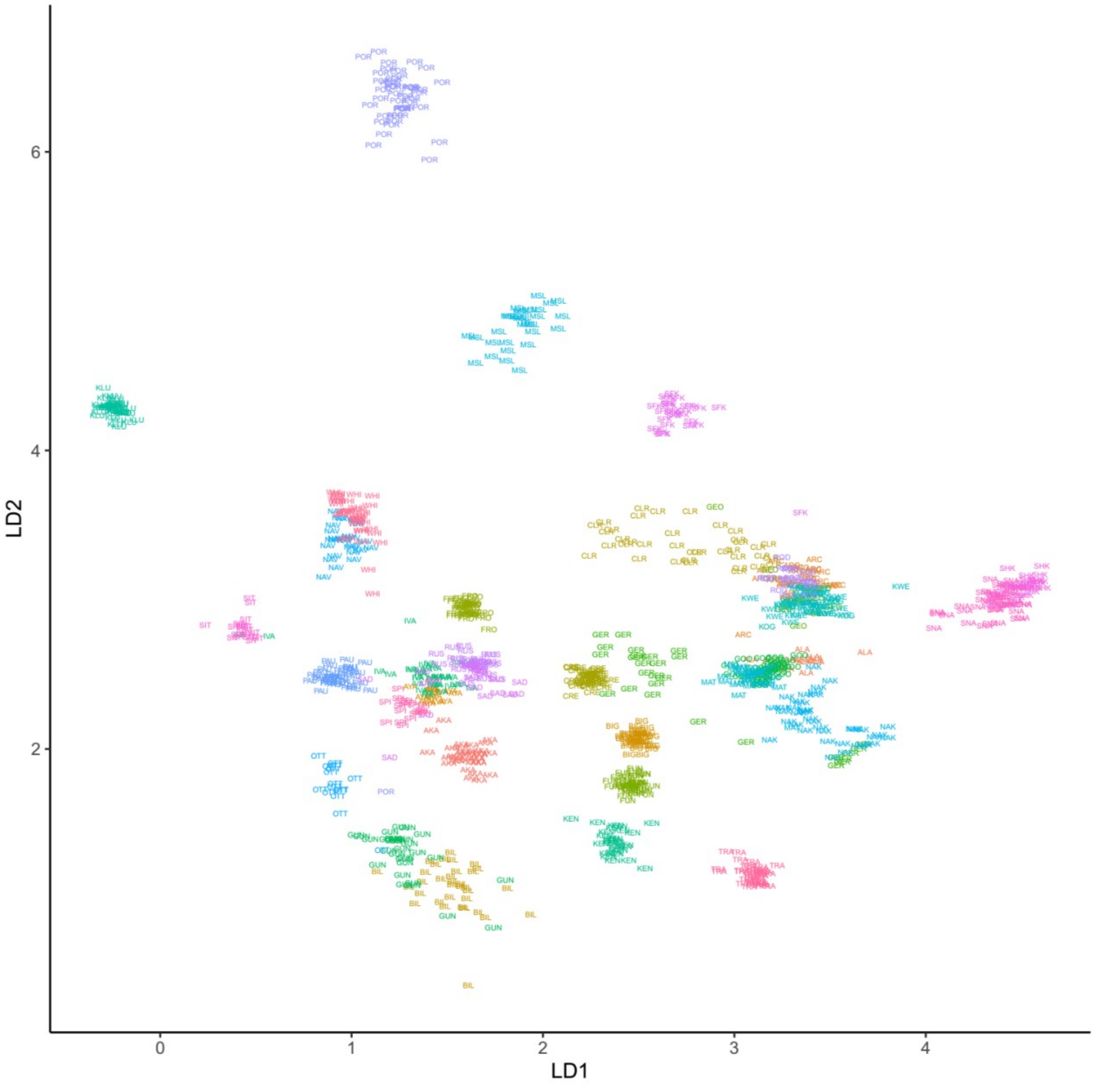


**E) Alaska**

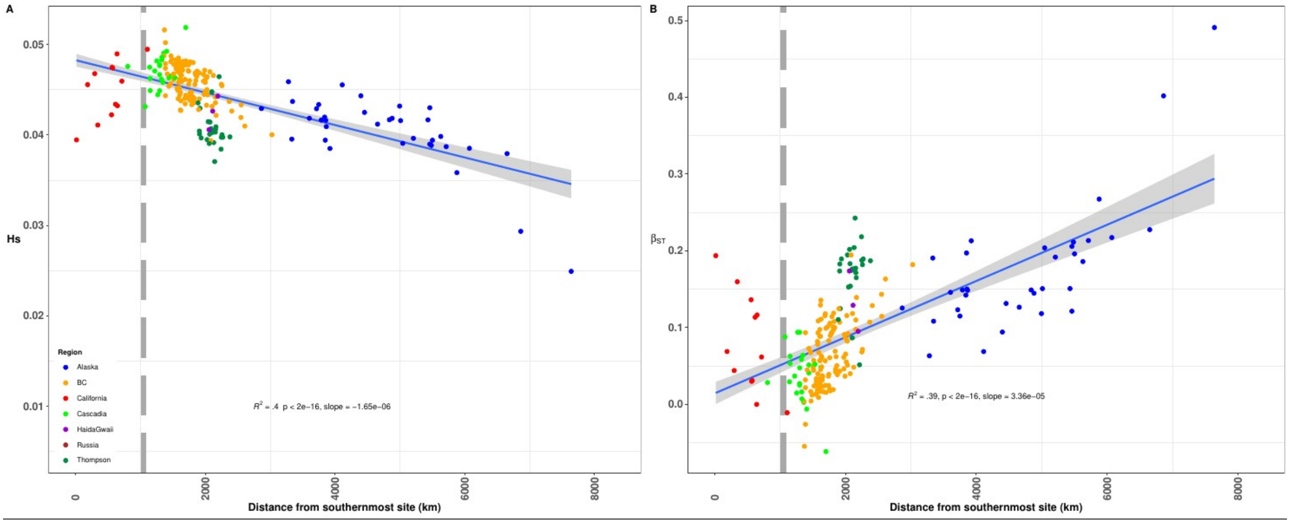


**FigureS06: Genetic diversity and differentiation.** A) Linear relationship between genetic diversity (Hs) and distance from the southernmost population located in California. B) Linear increase in genetic differentiation as measured by βST as a function of the distance from the southernmost population located in California. Negative values indicate the most likely ancestral population. The grey vertical bar in panels A and B indicates the approximate location of the southern limit of the icesheet at the end of the last glacial maximum. The grey shaded area along the regression line corresponds to the 95% confidence level interval obtained from the linear models applied to each dataset.

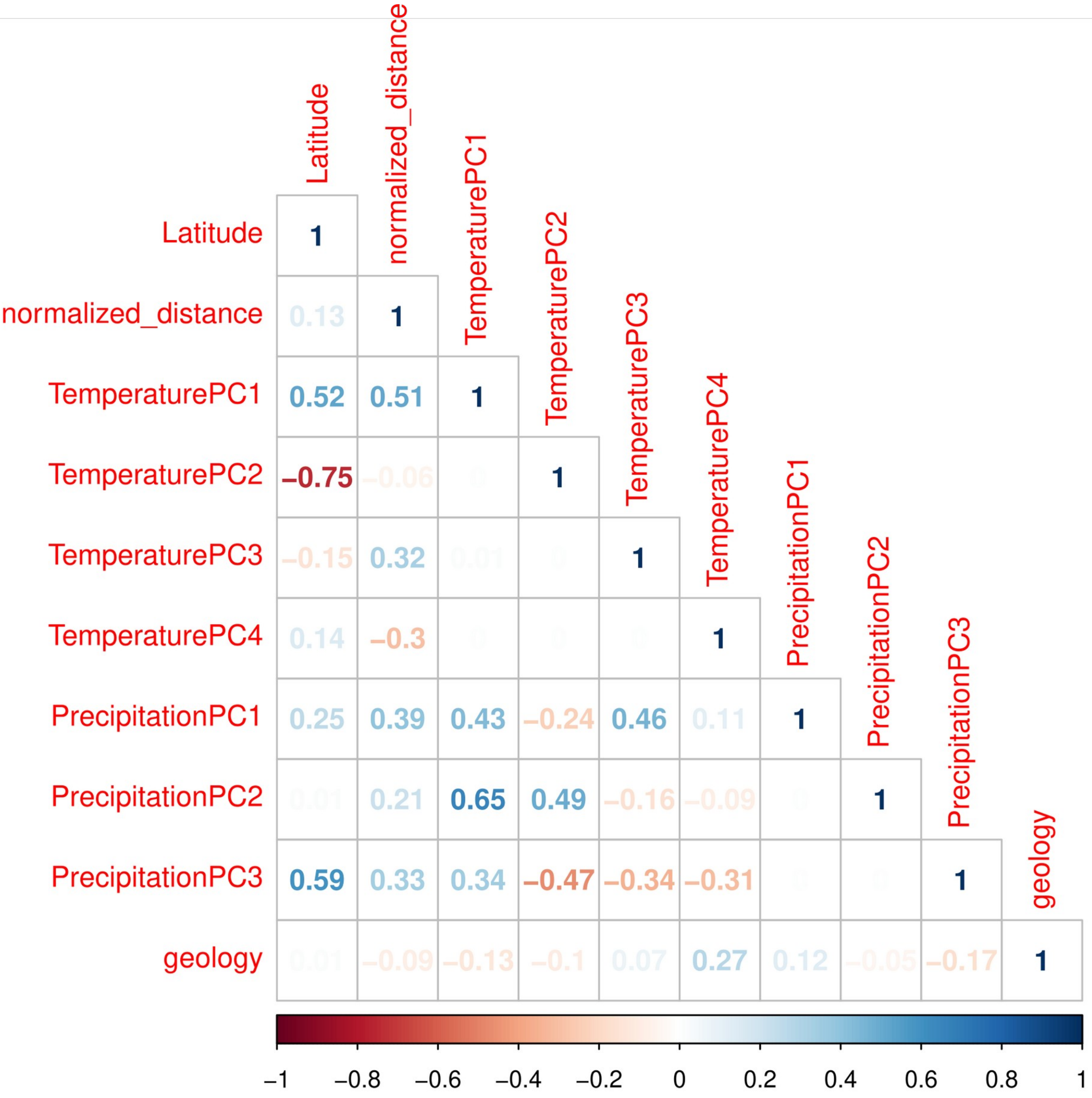


**Figure S07: Correlation plot among all variable included in the anaylses.** Empty cases correspond to value of 0.

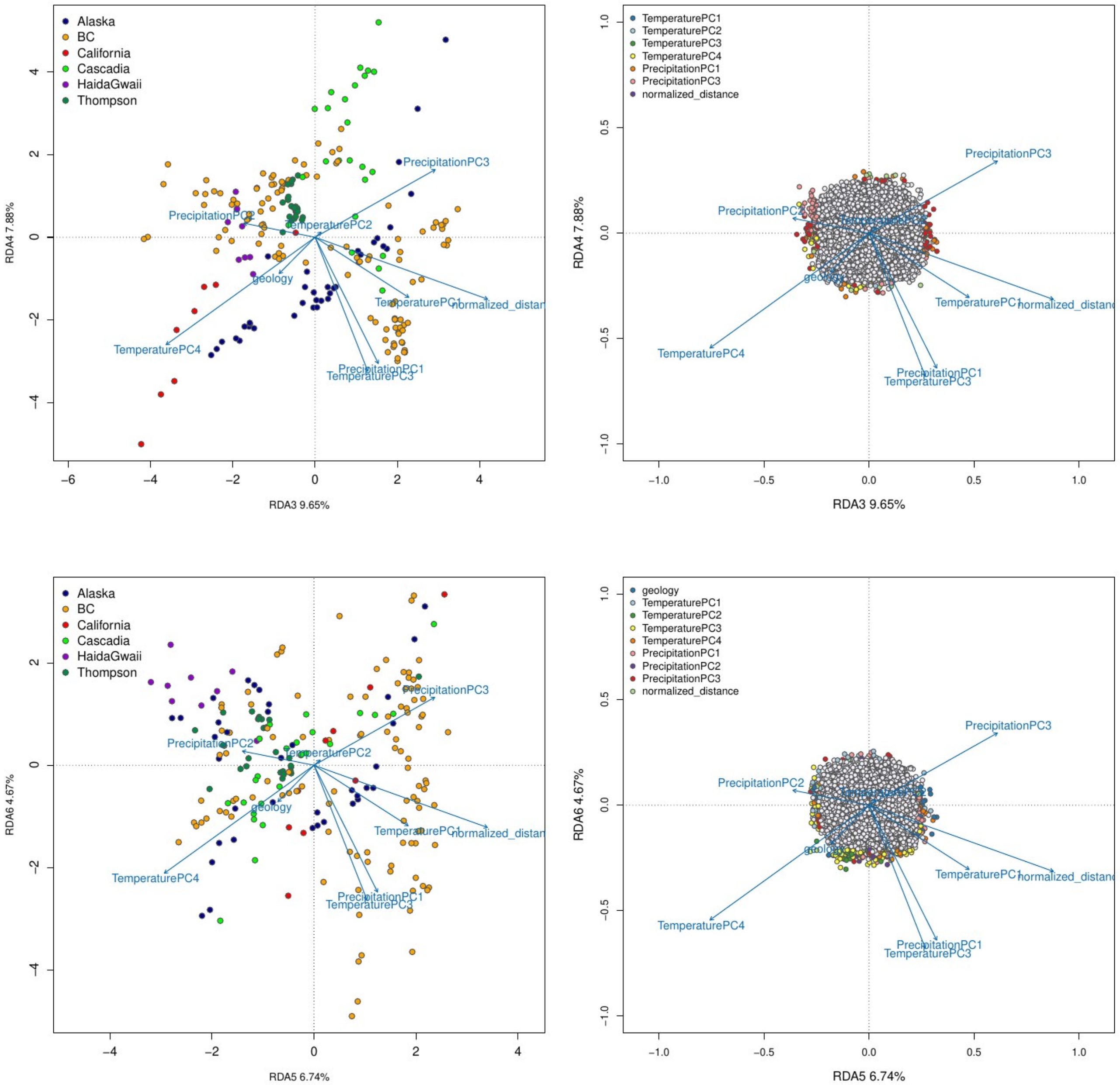


**Figure S08: RDA results along additional significant axes.** The top panels show the discrimination of populations along with environmental variables on axes 3-6; points are individuals colored according to the population from which they were sampled. The bottom panels show the outlier SNPs (colored points) associated with a given environmental variable on axes 3-6. Grey points correspond to non-outlier SNPs.

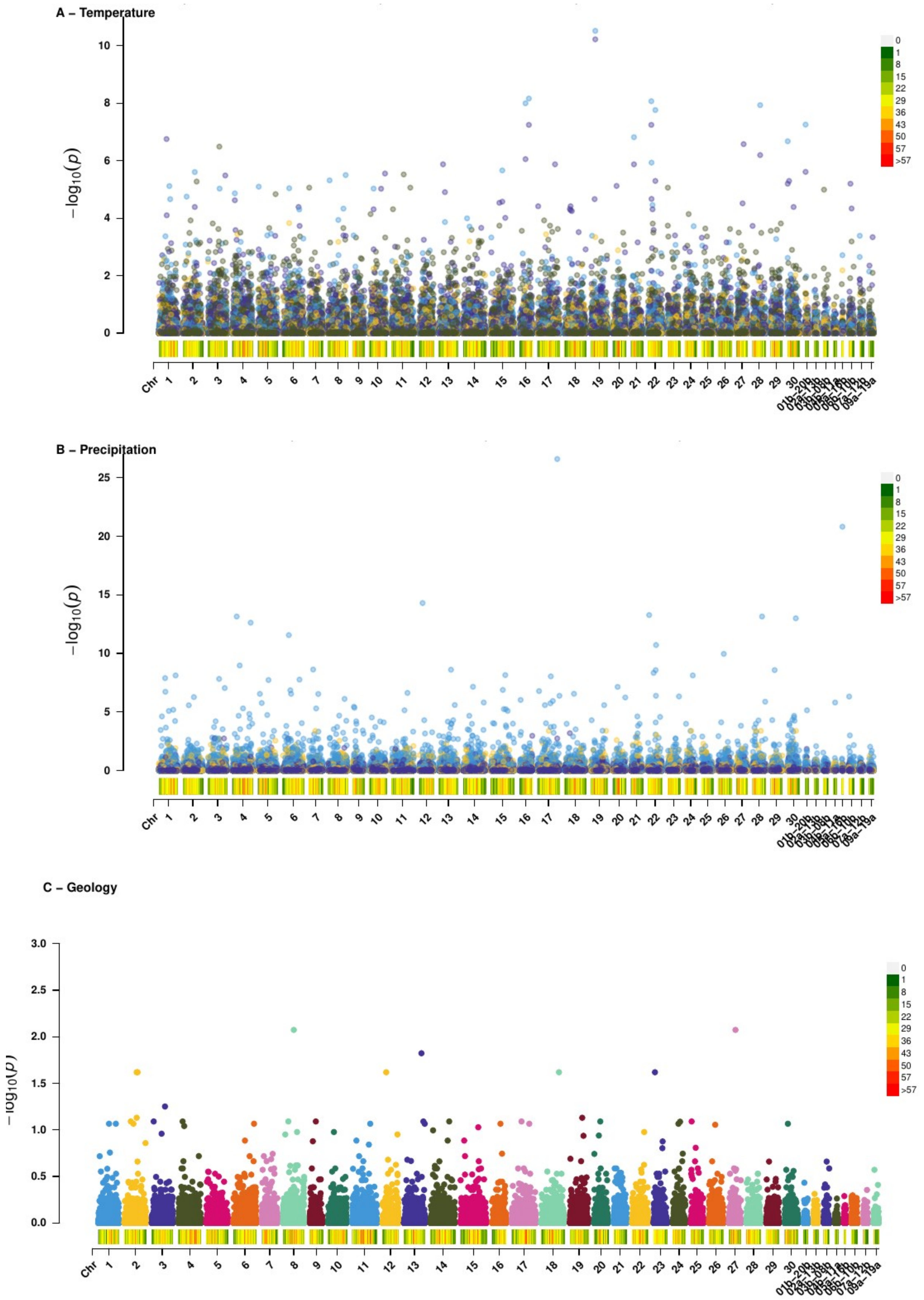


**Figure S09:** LFMM results for each bioclimatic variable (Temperature and Precipitation, reduced to three significant PC axis) and for one geological variable (Rock). x-axis = chromosome ordered from 1 - 30 (diploid) and chromosome with residual tetraploidy (from 01b_20b to 09a_19a). y-axis = -log(10) pvalue for each environmental variable tested.

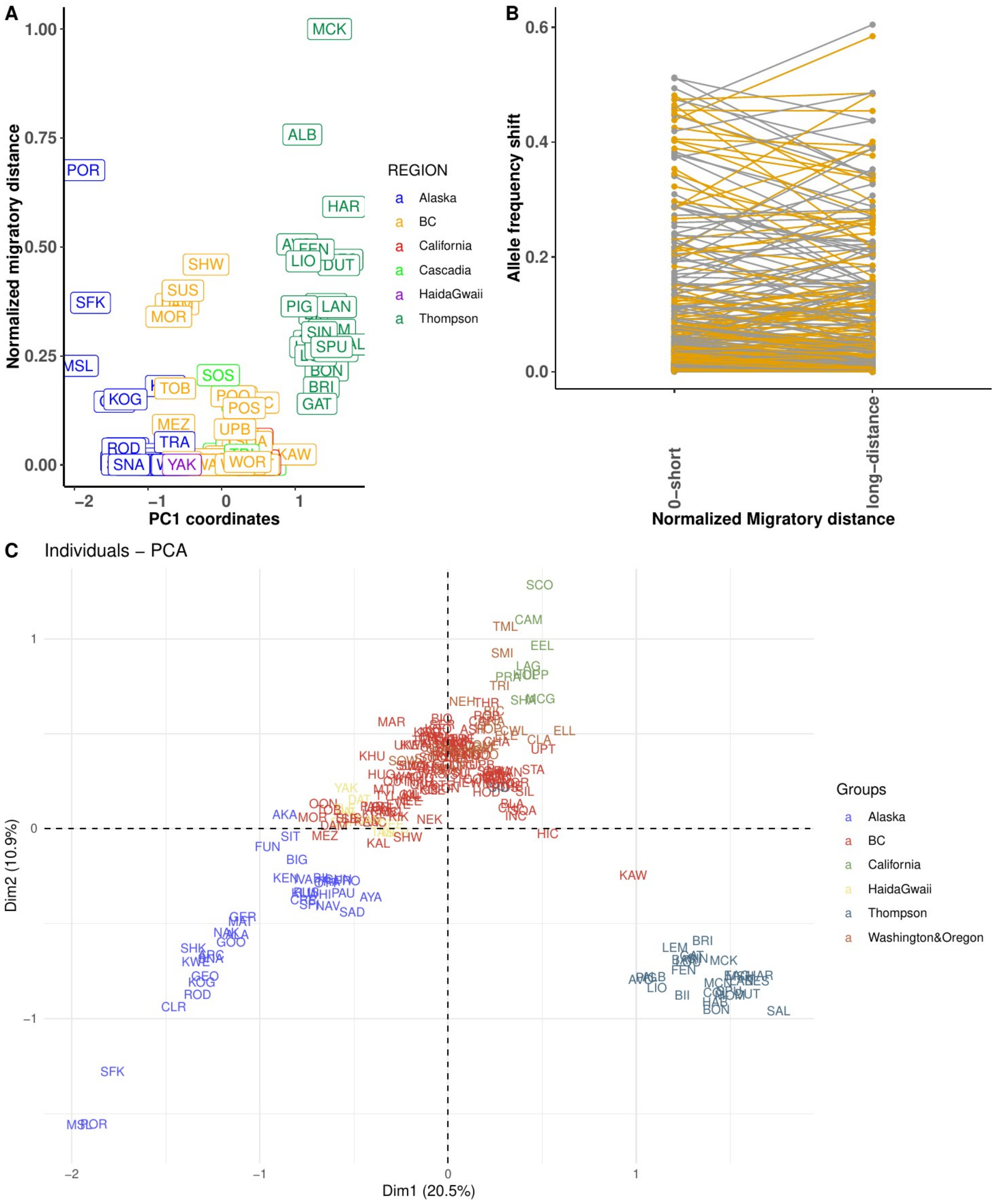


**Figure S10:** A) Discrimination along the PC1 axis between samples using a random set of SNPs B) Allele frequency shift associated with a random set of SNPs, C) PCA discriminating all the populations with the same set of random SNPs, reflecting global population structure. Here the first two axes were significant, whereas only the first axis was significant in A).

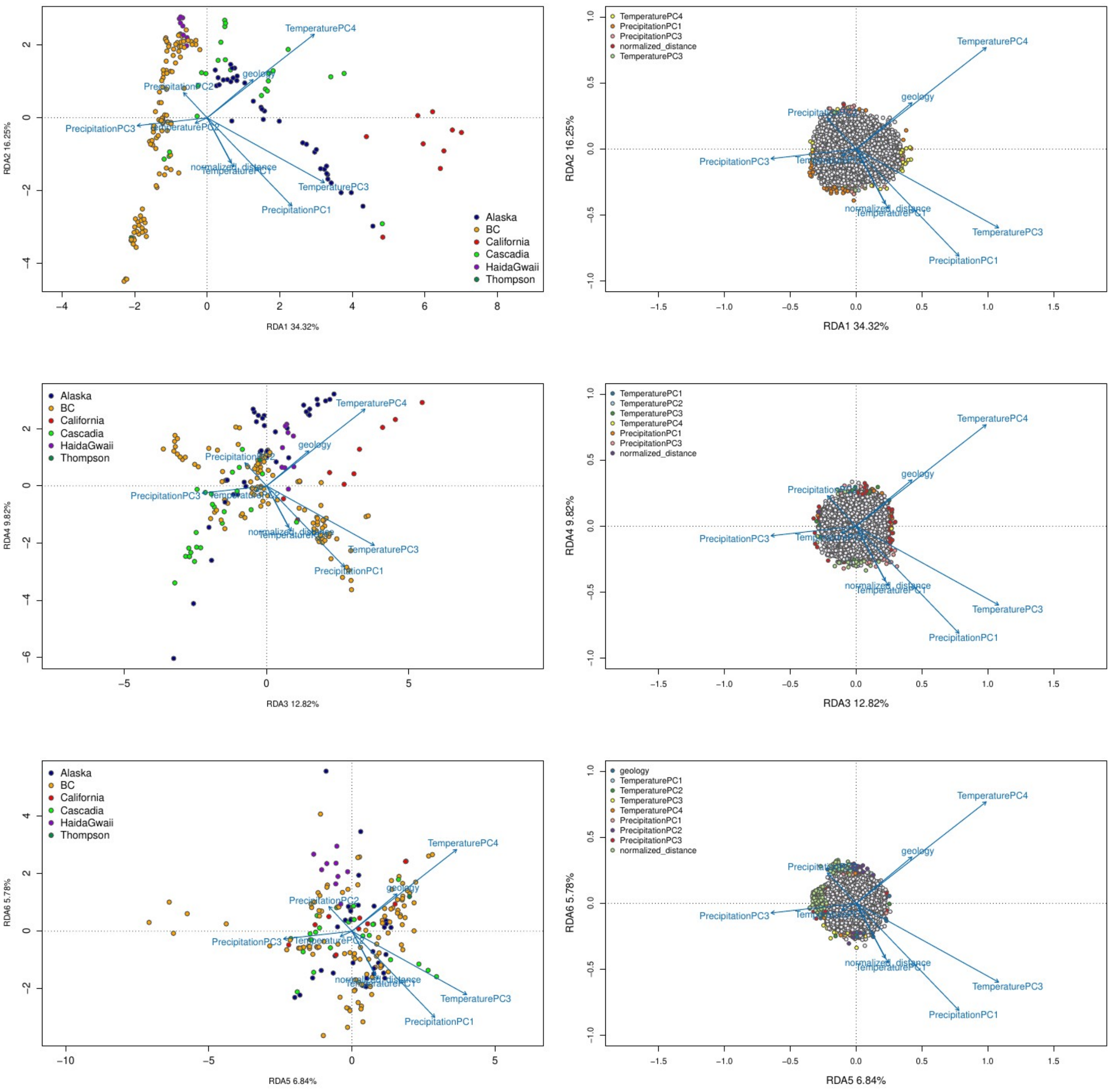


**Figure S11: RDA without Thompson Sites.** The top two panels show the discrimination of populations along with environmental variables on axes 1-2; points are individuals colored according to the population from which they were sampled. The middle two panels show the outlier SNPs (colored points) associated with a given environmental variable on axes 3-4. The lower panel gives the results for axis 5 and 6. Grey points correspond to non-outlier SNPs.

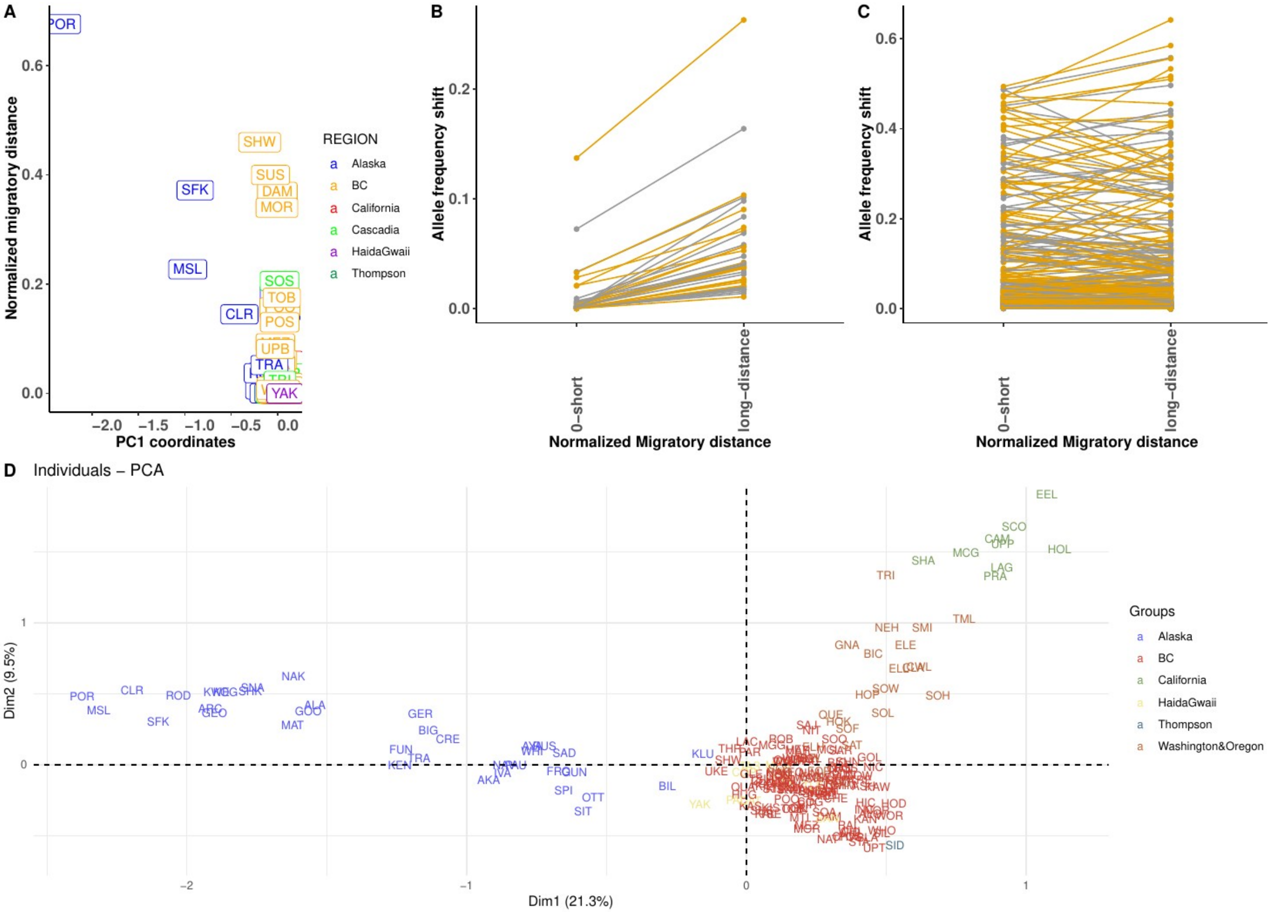


**Figure S12: Same as Figure 04 (maint text) and Figure S10 but without Thompson river.** A) Discrimination along the PC1 axis between samples associated with the set of shared outliers between the RDA and LFMM. B) Allele frequency shift associated with the set of outlier SNPs, C) Allele frequency shift when considering a set of randomly selected non-outlier SNPs and D) associated discrimination of the populations. Here the first two axes were significant, whereas only the first axis was significant in A).

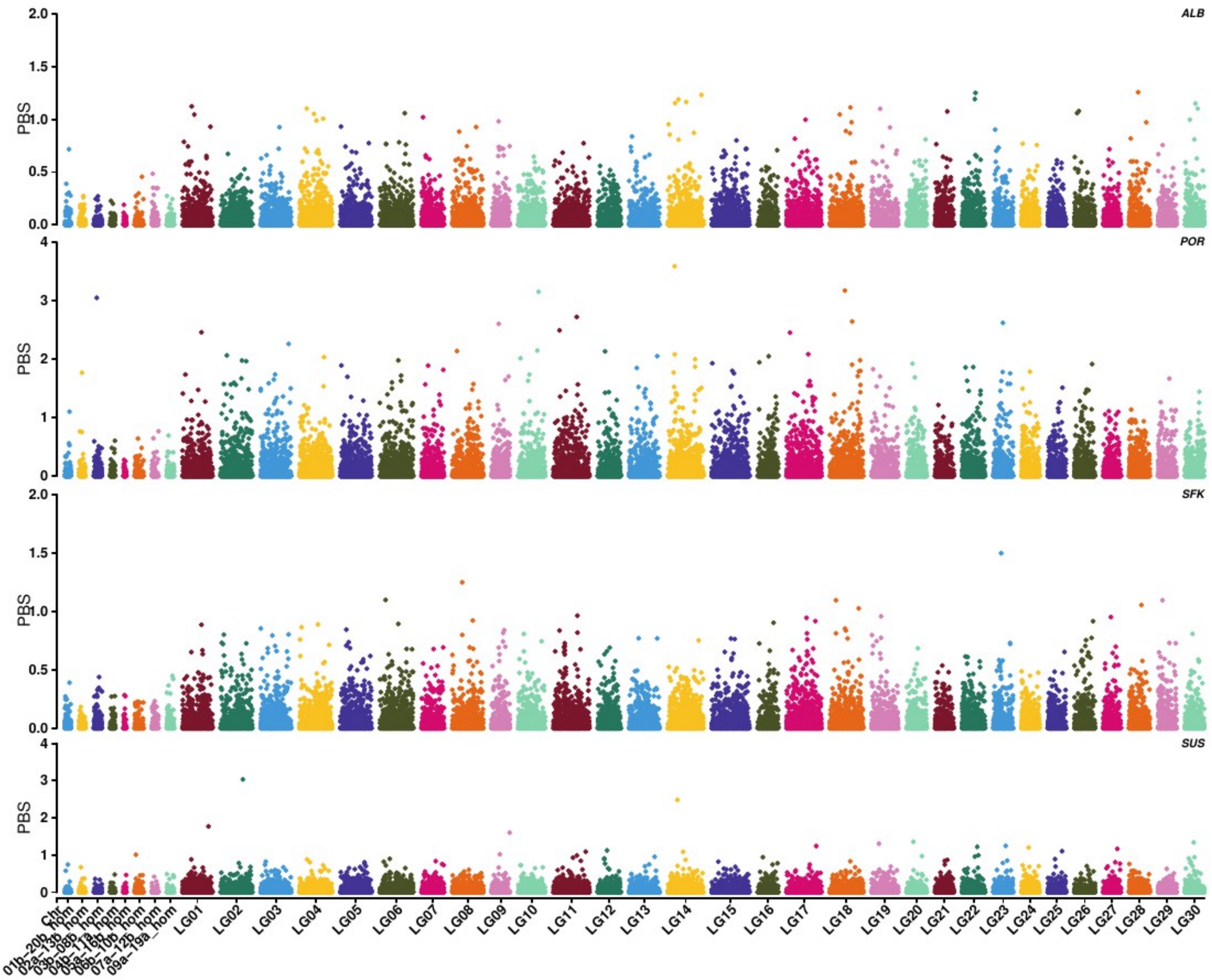


**Figure S13: PBS along the genome for multiple pairwise comparisons. Few peaks stands out and are shared among all populations.** TopRight initials provide the name of the upstream population. Each upstream population was compared to its downstream counterpart.

ALB = Albreda River (Upstream Thompson population), MAD = Mad River (Upstream Oregon population) POR = Porcupine River (Upstream Alaskan population), SFK = SFKuskokwim River (Upstream Alaskan population) SUS = Sustut River (Upstream BC population). The first 8 pairs of chromosomes corresponds to chromosomes with residual tetraploidy. LG01 to LG30 represents diploid chromosomes.

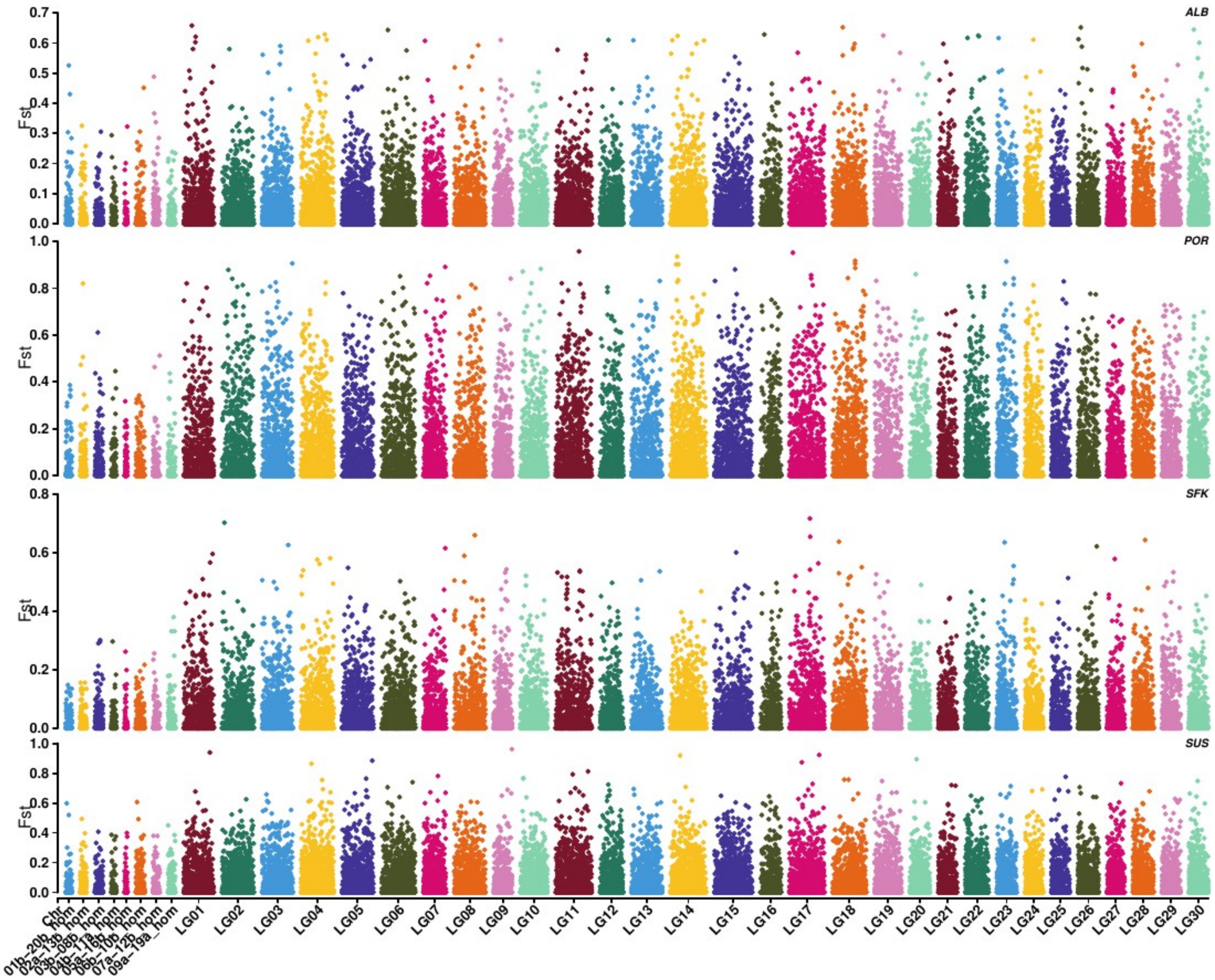


**Figure S14: Fst along the genome for multiple pairwise comparisons.** Each upstream population was compared to its downstream counterpart. TopRight initials provide the name of the upstream population. Each upstream population was compared to its downstream counterpart. ALB = Albreda River (Upstream Thompson population), MAD = Mad River (Upstream Oregon population) POR = Porcupine River (Upstream Alaskan population), SFK = SFKuskokwim River (Upstream Alaskan population) SUS = Sustut River (Upstream BC population). The first 8 pairs of chromosomes corresponds to chromosomes with residual tetraploidy. LG01 to LG30 represents diploid chromosomes.

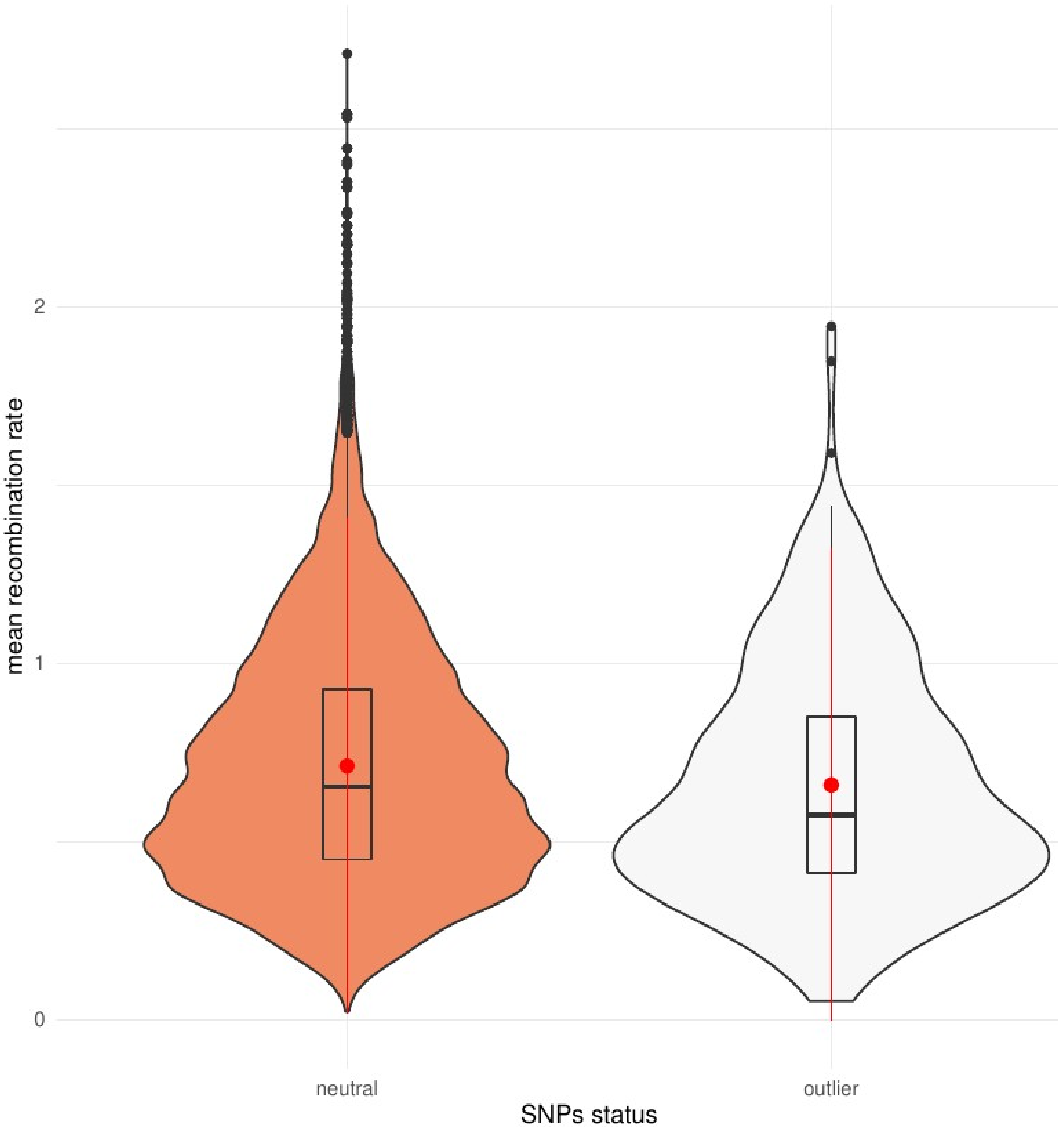


**Figure S15: Comparison of genome-wide populations scale recombination rate (rho = 4*Ne*4) for outlier vs non-outlier windows.**

**Figure S16: Map of genotypic distribution for outliers strongly correlated with environmental variation.**

**Outliers associated to distance:**
**SNP_ID:** 3912538:27:-
**Annotation:** Modifier, **Protein:** SH3 and multiple ankyrin repeat domains protein 2

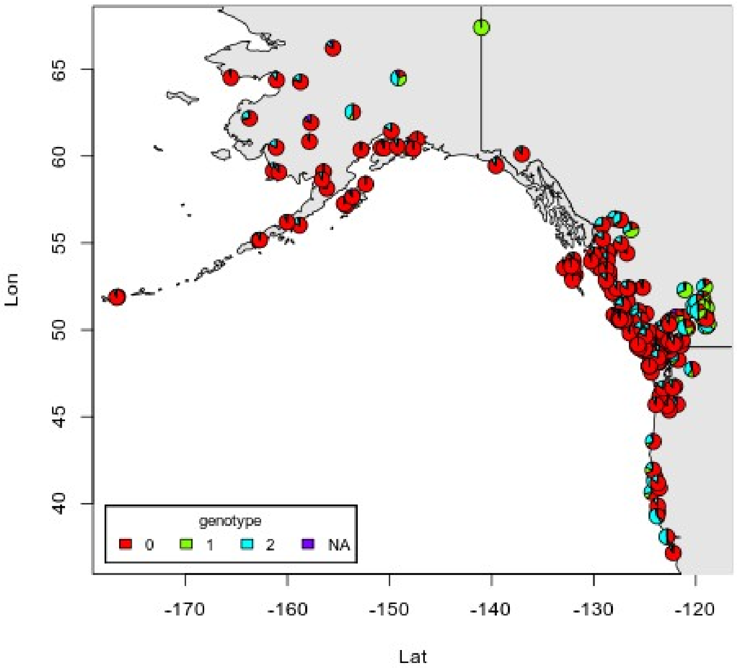


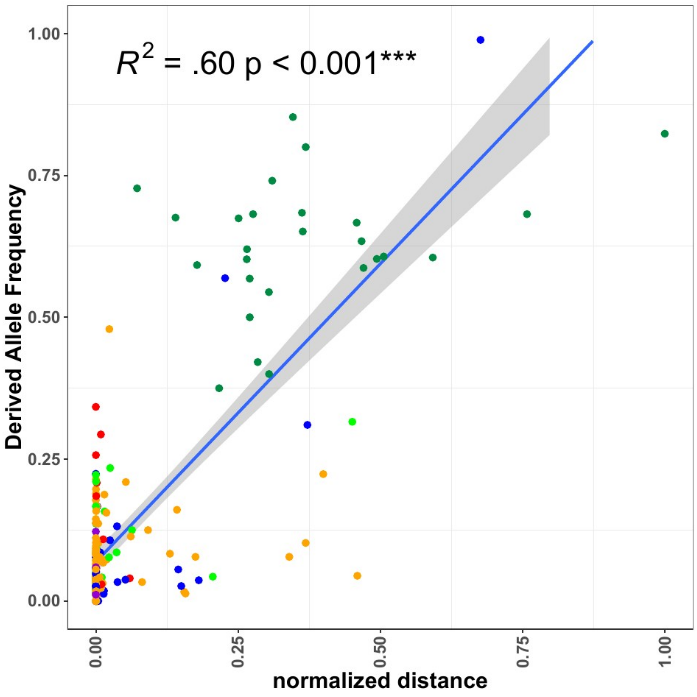


**SNP_ID: 4244396:56:+**
**Annotation: Intergenic**
**Protein: Epas1 – Pcgf5**

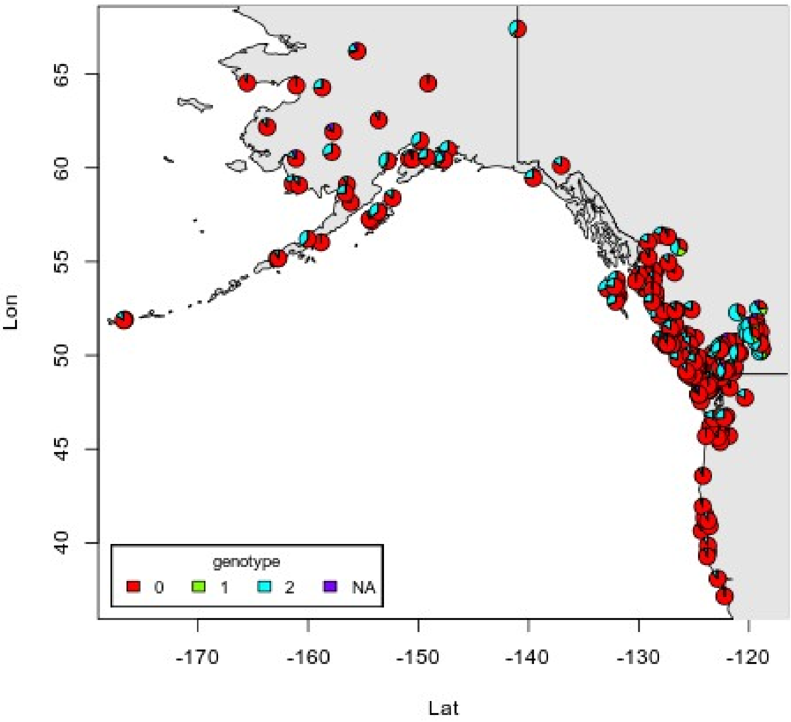


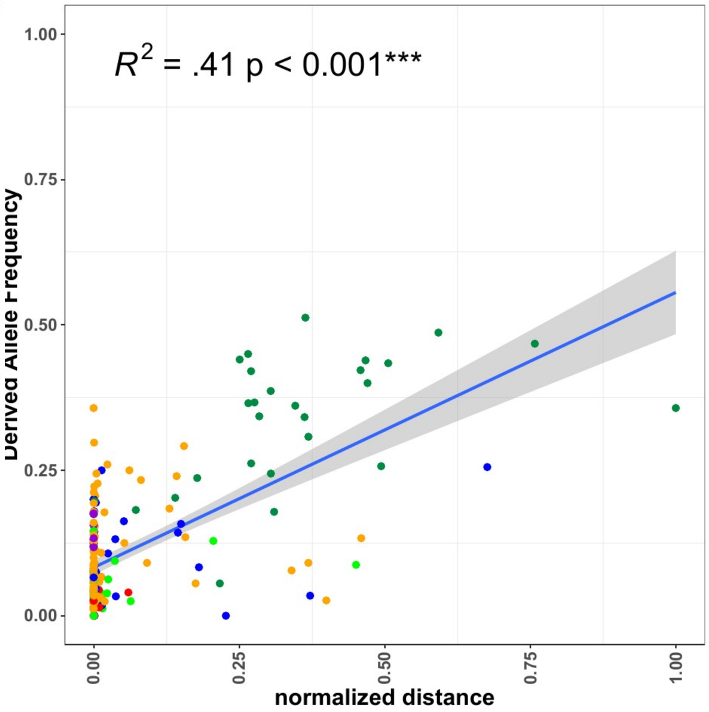

**Temperature Associated outliers:**

SNP_ID: 1031950:14:+
Location: Missense Variant – tankyrase
Linear Model MAF ~ Temperature 1 :
*pvalue* = 4e-26, **$R^2$ = 0.405**

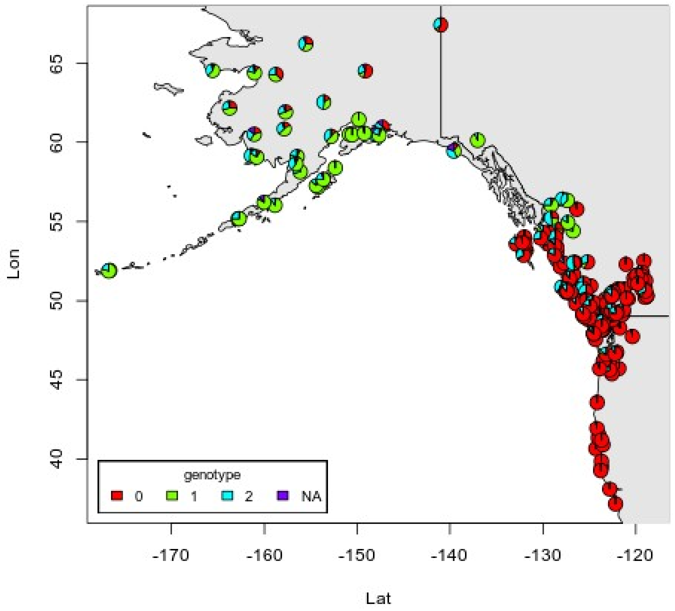


SNP_ID : 940737:12:+
Location: Intron – Kappa variable 6D—41
Linear Model MAF ~ Temperature 1 :
*pvalue* = 9.5e-26, **$R^2$ = 0.401**

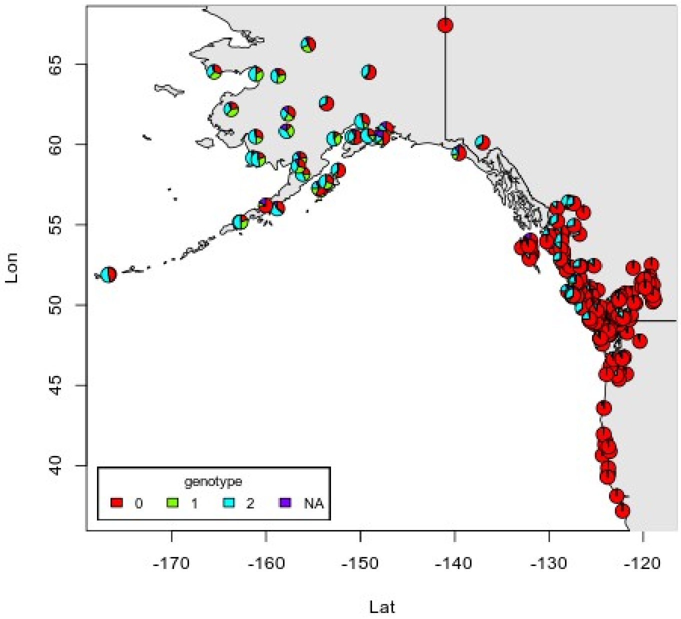


**Precipitation Associated outliers**

SNP_ID: 338867:44:-
Location: intron – interleukin-1 receptor type 1
Linear Model: (MAF ~ Precipitation):
*pvalue = 1.5e-24*, **R2 = 0.386**

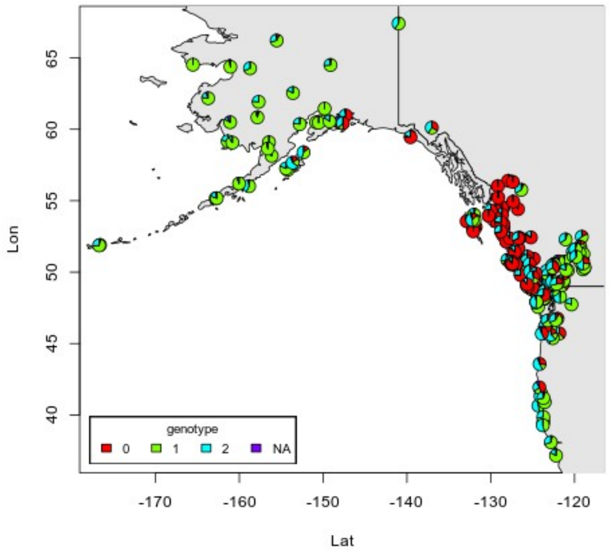


SNP_ID: 4591829:8:-
Location: intron – lrwd1
Linear Model (MAF ~ Precipitation1)
*pvalue = 5.7e-24*, **R2 = 0.378**

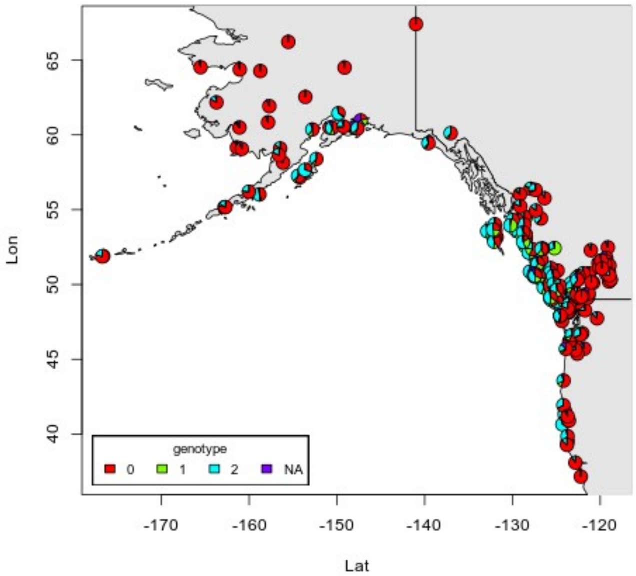
